# Proteomic signatures of cognitive resilience in LOU/c/Jall rats converge with inverse hippocampal axes of Alzheimer’s disease

**DOI:** 10.64898/2026.06.30.735140

**Authors:** L Gephine, AM Badina, S Corvaisier, BB Tournier, M Leger, T Freret

**Author notes:** Equal contribution (L Gephine, AM Badina).

## Abstract

Why some individuals maintain good level of cognitive performances during aging, others don’t or even progress toward Alzheimer’s disease. We profiled the hippocampal proteome of adult LOU/c/Jall rats, a strain associated with spontaneous cognitive longevity, and compared this proteomic state with a published human hippocampal Alzheimer’s disease dataset. Because individual protein changes did not survive proteome-wide correction, interpretation was based on convergent pathway-level, cell-type enrichment and cross-species directional analyses. The LOU hippocampus displayed a structured remodeling of mitochondrial, lysosomal, proteostatic and synaptic systems. Oligodendrocyte-associated nuclear-encoded complex I/III components were reduced, whereas neuronal mitochondrial aminoacyl-tRNA synthetases, V-ATPase, SNARE-related proteins and inhibitory-transmission markers were increased. CD200 was markedly reduced, but this occurred without accompanying complement, microglial, astrocytic or inflammatory activation signatures. Cross-species overlay indicated that several LOU-associated axes were directionally opposed to late Alzheimer’s disease, particularly synaptic vesicle and inhibitory-transmission programs, whereas myelin-associated changes occupied a lower-amplitude and non-inflammatory position along an axis altered in early Alzheimer’s disease. These findings identify a hippocampal proteomic configuration associated with the LOU resilience phenotype and suggest that successful brain aging and Alzheimer’s disease may involve opposing states of shared hippocampal molecular systems.

**GRAPHICAL ABSTRACT:** 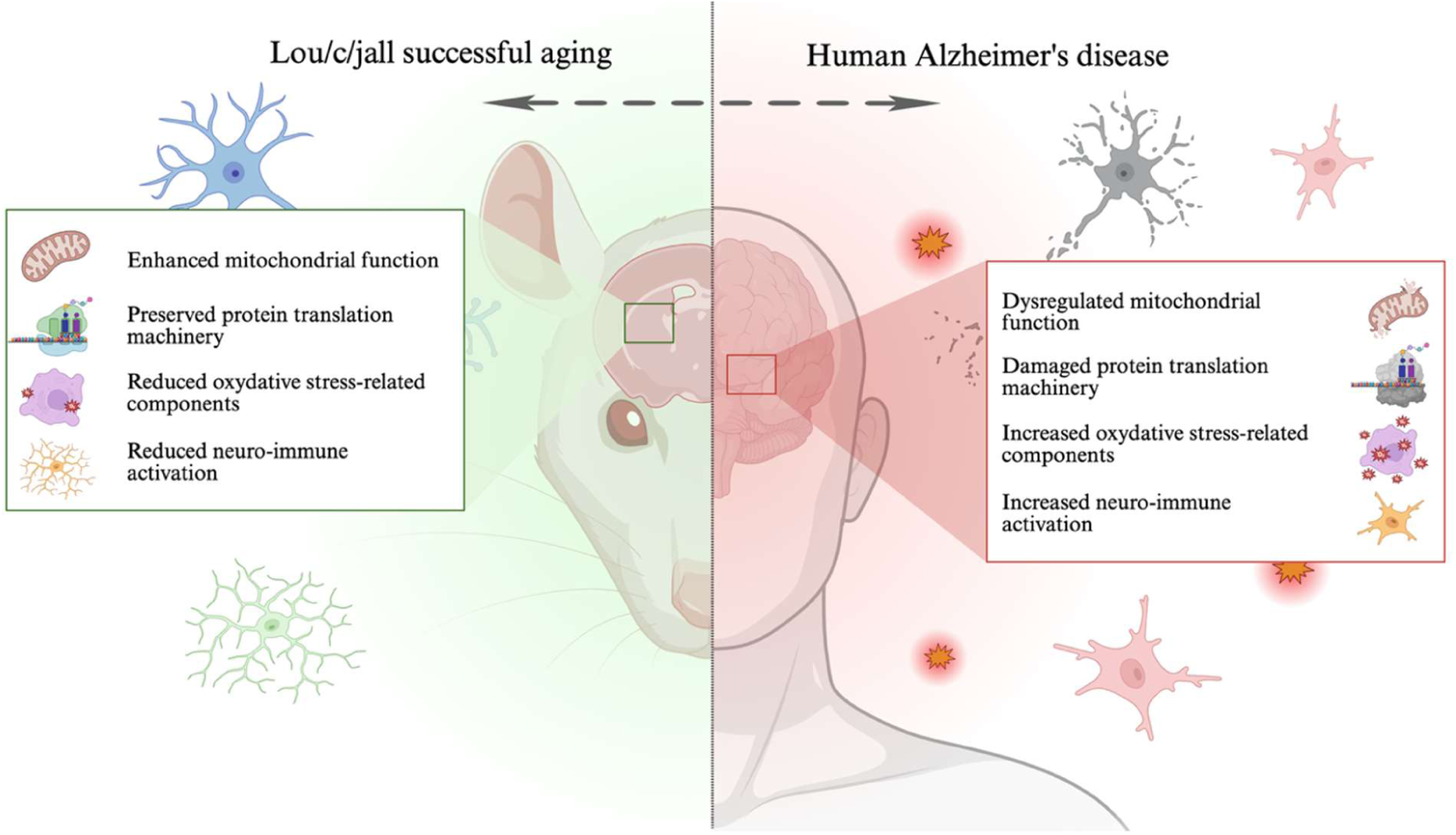

**HIGHLIGHTS:**

- Hippocampal proteome of the LOU/c/Jall rat at 3 months profiled by DIA-MS
- Coordinated reduction of complex I/III subunits in oligodendrocytes
- Neuronal aminoacyl-tRNA synthetases, V-ATPase and SNARE machinery up-regulated
- Marked reduction of CD200 with no inflammatory correlate
- Late human AD hippocampal transcriptome moves opposite to adult LOU

## 1. INTRODUCTION

Aging is a multifactorial biological process characterized by a progressive cognitive decline, even in the absence of pathology. In the brain, age-related changes manifest most consistently as impairments in learning, memory and executive function (Harada et al., 2013). However, certain individuals retain preserved cognitive function into advanced age, a phenomenon referred to as successful aging (Rowe & Kahn, 1997). Understanding the biological mechanisms that underlie successful aging is a central question of the field and could inform strategies to support brain health throughout the lifespan. Successful global aging has been linked to a combination of genetic, behavioral and environmental factors, including sustained physical activity, metabolic regulation and psychosocial engagement (Nyberg et al., 2012). At the cellular level, preserved mitochondrial function (Sun et al., 2016), low oxidative stress (Lopez-Otin et al., 2013), reduced neuroinflammation (Barrientos et al., 2015) and maintenance of synaptic plasticity (Burke & Barnes, 2006) have emerged as core hallmarks. However, inter-individual variability and limited access to postmortem brain tissue complicate mechanistic studies in humans, thereby highlighting the value of animal models for identifying mechanisms predictive of successful aging and for understanding processes preserved throughout the lifespan (Lopez-Otin et al., 2013; Nyberg et al., 2012).

The LOU/c/Jall (LOU) rat is a rodent strain that models successful aging. Derived from the Wistar (WIS) lineage, LOU rats exhibit a median lifespan of approximately 36 months, compared to about 24 months in WIS rats (Menard et al., 2015). The strain shows a low incidence of spontaneous age-related disease and metabolic stability into old age (Alliot et al., 2002). They show reduced adiposity, sustained insulin sensitivity and high locomotor activity in old age (Alliot et al., 2002; Bardag-Gorce et al., 1999; Veyrat-Durebex et al., 2009). Further, LOU rats retain intact cognitive performance well beyond the age at which other strains, such as WIS or Sprague-Dawley rats, display significant decline (Kollen et al., 2010; Menard & Quirion, 2012; Paban et al., 2013). While WIS and Sprague-Dawley rats show impairments in spatial learning and memory from 18-20 months of age, LOU rats maintain stable performance at 24-30 months, with no evidence of cognitive deterioration. These features make the LOU rat a compelling model for identifying endogenous protective mechanisms. Compared with WIS rats, adult LOU rats have been shown to exhibit enhanced mitochondrial efficiency and reduced oxidative damage in skeletal muscle and liver (Garait et al., 2005; Piec et al., 2005). In the hippocampus, transcriptomic studies across aging further identified differential expression of genes involved in lipid metabolism, nuclear architecture, immune regulation and inflammation in LOU rats relative to WIS rats (Paban et al., 2013). Finally, using an experimental model of sporadic Alzheimer’s disease, we recently demonstrated that LOU rats exhibit better cognitive performance than WIS rats despite a comparable cerebral amyloid burden, further supporting the LOU strain as a model of spontaneous cognitive resilience (Gephine et al., 2025).

The early hippocampal proteomic landscape of the LOU rat remains unexplored, although it may already reveal molecular adaptations that contribute to long-term cognitive preservation. This question is particularly relevant because, at 3 months of age, LOU rats already display molecular differences that are associated with preserved cognitive performance after STZ injection compared with WIS rats (Gephine et al., 2025). In the present study, we therefore first defined the hippocampal proteomic signature of successful aging by directly comparing adult LOU rats with age-matched WIS rats. As a second level of analysis, we analyzed the cortical proteome of a control bench of rat at the same age (LOU *versus* WIS), in order to determine whether the LOU-associated molecular adaptations reflect a widespread brain phenotype or a region-specific specialization of the hippocampus. Finally, as a third level of analysis, we examined the relevance of this rat hippocampal signature to human brain aging by overlaying it onto hippocampal transcriptomic data obtained from aged individuals spanning Alzheimer’s disease progression. This cross-species comparison was motivated by the fact that postmortem hippocampal profiling in Alzheimer’s disease consistently reveals synaptic, mitochondrial and oligodendrocyte dysregulation alongside amyloid and tau pathology (Kim et al., 2024; Wang et al., 2024). In our previously published cohort of 30 human hippocampi spanning Braak stages 2 to 6, we characterized the temporal dynamics of neuroinflammatory and glial remodeling during Alzheimer’s disease progression (Badina et al., 2025). Early AD was dominated by oligodendrocyte and myelin pathway loss, together with microglial activation and peripheral immune infiltration, whereas late AD showed reduced microgliosis, partial myelin recovery and severe synaptic transcript loss. These data provided a framework to test whether the LOU hippocampal proteomic configuration occupies a protective or inverse position along molecular axes altered during human pathological aging.

Accordingly, this study had three sequential aims. First, we characterized the hippocampal proteome of young adult LOU rats by quantitative mass spectrometry, using WIS rats as the reference strain, and interpreted the resulting signature through pathway-level analyses, including Over-Representation Analysis (ORA), Gene-Set Enrichment Analysis (GSEA) and cell-type assignment by Expression-Weighted Cell-Type Enrichment (EWCE). Second, we compared the hippocampal and cortical proteomes at the same age to determine whether the LOU signature was regionally specialized or reflected a broader brain-wide phenotype. Third, we projected the LOU hippocampal proteomic signature onto our published human Alzheimer’s disease hippocampal transcriptomic dataset to determine whether molecular features associated with successful aging in the rat align with, diverge from, or oppose cellular dysregulations observed during human pathological aging.

## 2. MATERIALS AND METHODS

### 2.1 Animals and tissue

Hippocampal proteomics were performed on 3-month-old male LOU rats (n=4; CURB, Caen, France) and age-matched WIS rats (n=4; Janvier Labs, Le Genest-Saint-Isle, France). Animals were housed in transparent polycarbonate cages (Tecniplast, 59.8×38×20cm, 3 per cage) with nesting material and a cardboard tube. Housing conditions were 22±2°C, 55±10% relative humidity, 12:12h light/dark cycle (lights on 07:30). Food and water were available *ad libitum*. Procedures complied with EU Directive 2010/63 and the ARRIVE guidelines under authorization APAFIS#38885-2022101410092500-v1. Animals were anaesthetized (isoflurane 5% in O₂/N₂O 30/70%), euthanized by cervical dislocation and decapitated. Hippocampi and cortices were rapidly dissected, snap-frozen in liquid nitrogen and stored at −80°C until analysis. Cortical proteomic measurements (n=7 per rat strain) were obtained from previous study (Gephine et al., 2025).

### 2.2 Mass spectrometry and protein quantification

Trypsin/Lys-C digestion, nano-LC separation and data-independent acquisition on a timsTOF Pro instrument (Bruker Daltonics) followed our previously described workflow (Gephine et al., 2025). DIA-NN was used for spectral library matching, protein identification and label-free MaxLFQ quantification.

### 2.3 Differential expression, contaminant filtering and DIA outlier

Differential abundance was assessed with limma (R 4.3.2) using empirical-Bayes-moderated t-statistics on log₂-transformed protein intensities. The operational threshold was a moderated p-value<0.05 with |log₂FC|≥0.18 (1.2-fold) (Kammers et al., 2015; Tyanova et al., 2016). Benjamini-Hochberg q-values were computed across the full set of 7,984 quantified proteins. Because non-perfused hippocampal homogenates can carry residual blood and immunoglobulin proteins that distort directionality at extreme log₂FC, we removed entries matching globin (Hbb, Hba, Hbe, Hbq, Hbz, Mb, Cygb), immunoglobulin (Ig, Igh, Igk, Igl*, Jchain), or plasma/acute-phase carrier patterns (Alb, Ttr, Apoa1/2/4, Apob, Apoc1/2, Apoh, Apom, Trf, Hp, Hpx, Cp, Serpinc1, Serpina1/3, Fga, Fgb, Fgg, A2m, F2). Seven UniProt accessions BLASTed to globin or Ig families despite missing GeneSymbol annotation and were also excluded (A0A8L2R1Q6, F1LW33, A0A1K0H3R5, A0A8I6A9D7, A0A8I6AAM1, A0A0G2K828, A0A8I6AFR8). Thirty-five proteins were excluded in total.

Among the eleven mtDNA-encoded Electron Transport Chain (ETC) subunits quantified, mt-Nd2 was the only protein showing a large isolated decrease. Because DIA-MS estimates protein abundance from peptide- and fragment-ion-level evidence, proteins supported by few detected peptides are more vulnerable to unstable quantification. The mt-Nd2 signal had low peptide coverage and was not supported by a coordinated decrease across the other mtDNA-encoded ETC subunits. It was therefore retained in the master table but excluded from figures and from the GSEA ranked vector.

### 2.4 Functional and cell-type enrichment

Over-representation analysis (ORA) was performed *via* gseapy.enrichr (Chen et al., 2013) against KEGG_2021_Human, GO Biological Process 2023, Reactome 2022, MSigDB Hallmark 2020 and WikiPathway 2023 Human. The cleaned full proteome (7,949 entries) was used as the background set. Rat gene symbols were converted to upper-cased to enable cross-species mapping to human-curated libraries.

Gene-Set Enrichment Analysis (GSEA) was performed on the full proteome with gseapy.prerank. The ranking metric was defined as −log₁₀(p) × sign(log₂FC), thereby integrating both statistical evidence and direction of regulation. Analyses were run with 2,000 permutations and restricted to gene sets containing 5–500 proteins. Reported q-values correspond to Benjamini–Hochberg-adjusted p-values calculated within each library.

Expression-Weighted Cell-type Enrichment (EWCE) was performed in R with the ewceData mouse hippocampal single-cell reference dataset at annotation level 1 (Skene & Grant, 2016). Rat proteins were maps to mouse ortholog using g:Profiler (Raudvere et al., 2019), and enrichment was assessed with 10,000 bootstrap iterations against the cleaned proteome background. EWCE driver-gene lists were subsequently annotated for pathway enrichment using clusterProfiler and ReactomePA; the top 200 drivers were used for neuronal populations, whereas the complete driver list was retained for oligodendrocytes.

### 2.5 Cross-species comparison with human Alzheimer’s disease

We re-used differential expression tables from our published 770-gene neuroinflammation hippocampal cohort (Badina et al., 2025): 6 controls (Braak 2-3, CT), 12 early-AD (Braak 4, EAD) and 12 late-AD (Braak 5-6, LAD) subjects, with EAD-vs-CT and LAD-vs-CT DEG tables. Both datasets were restricted to the 757 protein-coding endogenous genes on the panel; symbol matching used the uppercase rat-to-human form. Direction concordance was tested with the binomial sign test against the 0.5 null, and quantitative agreement with Spearman rank correlation. Per-axis sign and rho statistics were computed on curated marker sets covering vesicular synaptic, inhibitory synaptic, general synaptic, oligodendrocyte/myelin, OPC, astrocytic, microglial/complement, mitochondrial OXPHOS, neuro-immune (CD200/CD200R1) and stress/proteostasis categories. ORA on the EAD and LAD DEG lists used the panel as background, otherwise identical settings to the LOU side. For Fig. 5B-C, the neuronal/synaptic, homeostatic microglia, DAM and central/peripheral immunity modules were additionally scored by GSVA (GSVA 2.2.1, Gaussian kernel; minSize=1 to retain small shared-gene modules) on the LOU/WIS log₂ imputed hippocampal protein matrix and on the published log₂ NanoString human matrix; LOU group effects used Welch two-sample t-tests, whereas human CT-LAD contrasts used one-way ANOVA followed by Tukey HSD. CD200 was shown separately as a direct protein/transcript contrast because it is a single-gene node.

With LOU and WIS rats (n=4 per group), no individual protein survives Benjamini-Hochberg correction at q<0.10. We therefore based inference and biological interpretation on pathway-level GSEA (within-set FDR), cell-type EWCE (permutation FDR) and the cross-species sign test (binomial test against the 0.5 null).

## 3. RESULTS

### 3.1 LOU *versus* WIS hippocampal proteomic landscape

A total of 7,984 hippocampal proteins were quantified across the two strains (n=4 per group). Before data cleaning, 481 proteins met the predefined nominal abundance-change criteria (p<0.05, |log₂FC|≥0.18). Eight of those proteins were flagged as blood- or immunoglobulin-derived contaminants. Notably, the three with the largest log₂FC values in the entire dataset (A0A8L2R1Q6, F1LW33 and Hbb, +4.5 to +5.9 in LOU) all belonged to the hemoglobin family. Because their inclusion would have artificially exaggerated the apparent directional imbalance between proteins increased and decreased in LOU rats, they were excluded from downstream analyses, together with the mt-Nd2 outlier described above (See section 2.3). After this filtering step, 473 proteins remained in the nominally altered protein set, with 308 increased and 165 decreased in LOU compared with WIS rats (Fig. 1).

**Figure 1.**
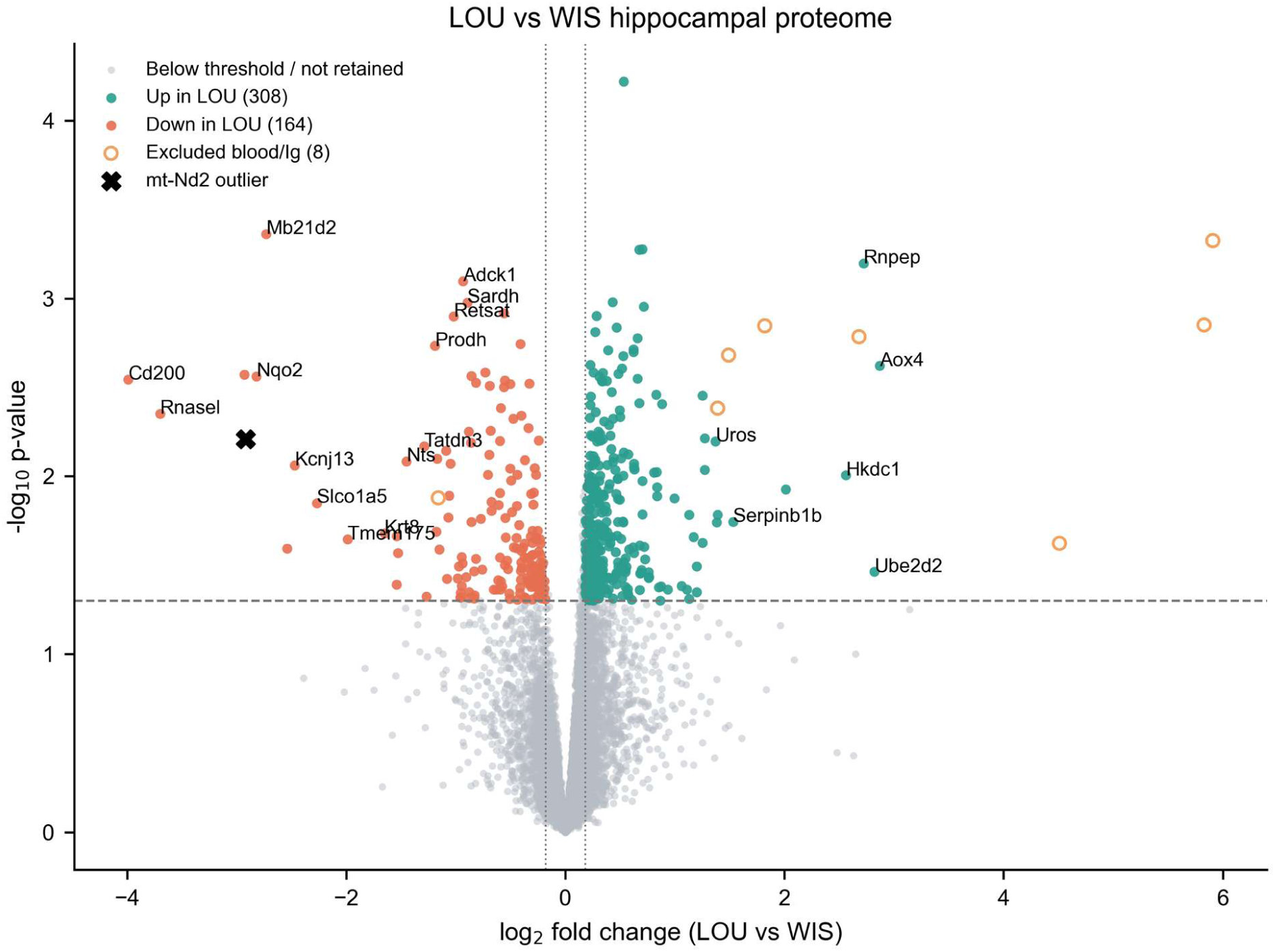
LOU *vs* WIS hippocampal proteomic landscape at 3 months of age. Volcano plot of log₂ fold change (LOU *versus* WIS) against −log₁₀ p-value from limma empirical-Bayes-moderated tests on n=4 per group hippocampi. Up-regulated retained proteins (308) are shown in green; down-regulated retained proteins (164) in red; below-threshold or non-retained proteins in grey. Open orange circles mark excluded blood or immunoglobulin contaminants that pass the operational thresholds (n=8; 35 contaminants excluded from downstream analyses in total). The top 25 proteins by combined log₂FC×−log₁₀p are labelled. Operational thresholds: p<0.05 and |log₂FC|≥0.18.

Except for two marked down-regulated proteins (namely Cd200 and Rnasel, log₂FC=−3.99 and −3.70, p=0.003 and 0.004), the differentially abundant proteins were distributed within a relatively modest log₂FC range, both for up- and down-regulated proteins, with median log₂FC values of +0.31 and −0.49, respectively. After multiple-testing correction, no individual protein reached significance using the Benjamini–Hochberg procedure at q<0.10, consistent with the limited statistical power expected in small-N DIA proteomics datasets (Tyanova et al., 2016). We therefore prioritized convergent evidence across complementary analytical frameworks rather than interpreting single-protein changes in isolation. These included pathway-level gene-set enrichment analyses against KEGG and MSigDB Hallmark collections, cell-type enrichment analysis using EWCE, and cross-species directional concordance analyses. Each analytical layer reached significance after its own multiple-testing correction (Sections 3.2–3.7), providing mutually reinforcing evidence for a coherent biological signal despite the absence of individually FDR-significant proteins.

### 3.2 LOU *versus* Wistar differentially abundant proteins pathway enrichment

ORA of the 473 differentially abundant proteins revealed a coherent functional partition between up- and down-regulated pathways (Fig. 2A-B), which was further supported by pre-ranked GSEA on the full proteome (7,984 proteins, Fig. 2C-D).

**Figure 2.**
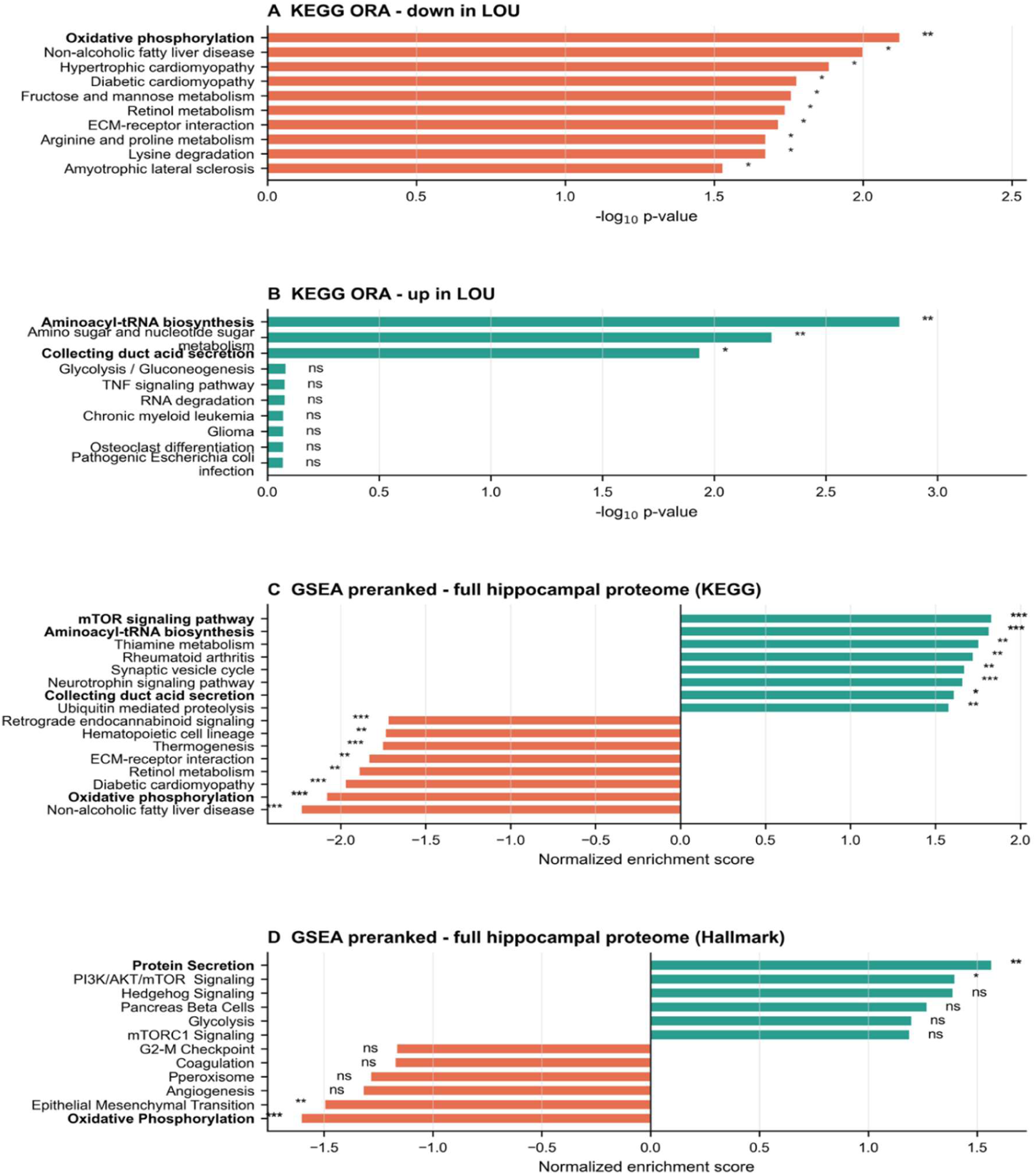
Pathway enrichment of LOU *versus* WIS hippocampal proteins. **A.** KEGG ORA on the 165 down-in-LOU proteins; the top hit is Oxidative Phosphorylation (six nuclear-encoded complex I/III subunits in the leading list). **B.** KEGG ORA on the 308 up-in-LOU proteins; the top hits are Aminoacyl-tRNA biosynthesis (predominantly mitochondrial ARSs), Amino sugar / nucleotide sugar metabolism, and the V-ATPase H⁺-transporting subunits (annotated by KEGG as Collecting duct acid secretion). **C.** Pre-ranked gene-set enrichment analysis (GSEA) on the full hippocampal proteome (rank score=−log₁₀(p)·sign(log₂FC)) against KEGG_2021_Human; bars show normalized enrichment scores for terms reaching FDR q<0.25. The mt-Nd2 outlier was excluded from the ranked vector. **D.** GSEA against MSigDB Hallmark sets; positive enrichment of Protein Secretion and PI3K/AKT/mTOR Signaling, and negative enrichment of Oxidative Phosphorylation, confirm the same partition independent of the pathway library. Bar labels show nominal p-value thresholds (* p<0.05; ** p<0.01; *** p<0.001; ns, p≥0.05).

Among the 165 down-regulated in LOU rats (Fig. 2A), ORA identified revealed enrichment in Oxidative Phosphorylation (KEGG, p=0.0075), driven by nuclear-encoded subunits of complex I and complex III genes (NDUFA9, NDUFA13, NDUFB10, NDUFA5, NDUFS1 and UQCRC2). Consistently, pre-ranked GSEA against the KEGG database confirmed a significant negative enrichment of Oxidative Phosphorylation across full proteome (NES=−2.08, FDR q=0.005, Fig. 2C), with 36 of 81 genes contributing to the leading edge. This leading edge included several nuclear-encoded complex I/III components, such as NDUFA13, NDUFA5, NDUFS1, UQCRC2, NDUFA9, NDUFB10, NDUFS4, NDUFS7, NDUFA6, NDUFV1 and NDUFA10. Of note, no mtDNA-encoded subunit was retained. Analysis using the independent MSigDB Hallmark library showed a concordant negative trend for Hallmark Oxidative Phosphorylation, although this did not reach conventional FDR significance (NES=−1.60, q=0.26) (Fig. 2D). Together, these results indicate that the down-regulated arm of the LOU hippocampal signature is dominated by a reduction in nuclear-encoded respiratory-chain components, particularly complex I and complex III subunits, rather than by a coordinated decrease in mtDNA-encoded ETC proteins.

Enrichment analysis of the 308 proteins increased in LOU rats (Fig. 2B) identified KEGG aminoacyl-tRNA biosynthesis as the top enriched pathway (p=0.0015), driven predominantly by mitochondrial aminoacyl-tRNA synthetase and related factors (including LARS2, MTFMT, TARS3, QRSL1, VARS2, YARS2 and CARS2). Another enriched KEGG term, collecting duct acid secretion, was driven by the V-ATPase H⁺-transporting subunits (including ATP6V1G2, ATP6V1E1 and ATP6V1C1) indicating that lysosomal functions and vesicular proton-pump components contributed to the up-regulated part of the LOU signature.

Pre-ranked GSEA on the full proteome confirmed the enrichment observed among up-regulated proteins. KEGG Aminoacyl-tRNA biosynthesis (Fig. 2C) reached significant positive enrichment (NES=+1.81, FDR q=0.036), with 27 of 42 genes contributing to the leading edge (including YARS2, TARS3, CARS2, LARS2, FARSB, MTFMT, VARS2, QRSL1, WARS2, HARS1, FARS2 and DARS2). Those genes pointing to a predominantly mitochondrial translation-related signature. KEGG mTOR signaling was also positively enriched (NES=+1.83, FDR q=0.049), with leading-edge genes including LAMTOR3, PTEN, ATP6V1E1, MIOS, ATP6V1C1, ATP6V1G2, DEPDC5, ATP6V1D, ATP6V1B1 and RPTOR. In the independent MSigDB Hallmark library (Fig. 2D), Hallmark Protein Secretion showed a concordant positive trend (NES=+1.56, q=0.18), driven in part by SNARE/vesicular machinery (such as STX7, VAMP7, NAPG, AP3S1, RER1 and VPS45).

Taken together with the negative enrichment of oxidative phosphorylation among down-regulated proteins, these results indicate that the LOU hippocampal proteomic signature comprises two complementary arms: a reduced nuclear-encoded respiratory-chain protein footprint, and a positive enrichment of mitochondrial aminoacyl-tRNA synthetases, V-ATPase/lysosomal proton-pump components, mTOR-pathway genes and vesicular secretion machinery. This pattern suggests a coordinated remodeling of mitochondrial, lysosomal and vesicular systems rather than a uniform increase or decrease in cellular energy metabolism.

### 3.3 Cell-type assignment

EWCE cleanly separated the LOU’s downregulated and upregulated proteins between glial and neuronal compartments (Fig. 3A–B). Down-regulated proteins enriched for oligodendrocytes (q=0.012; Fig. 3A). Up-regulated proteins enriched for neurons, including CA1 pyramidal neurons (q < 0.001), pyramidal neurons (q<0.001) and interneurons (q<0.001; Fig. 3B). No glial cell type was enriched in the up-regulated set, and no neuronal cell type in the down-regulated set.

**Figure 3.**
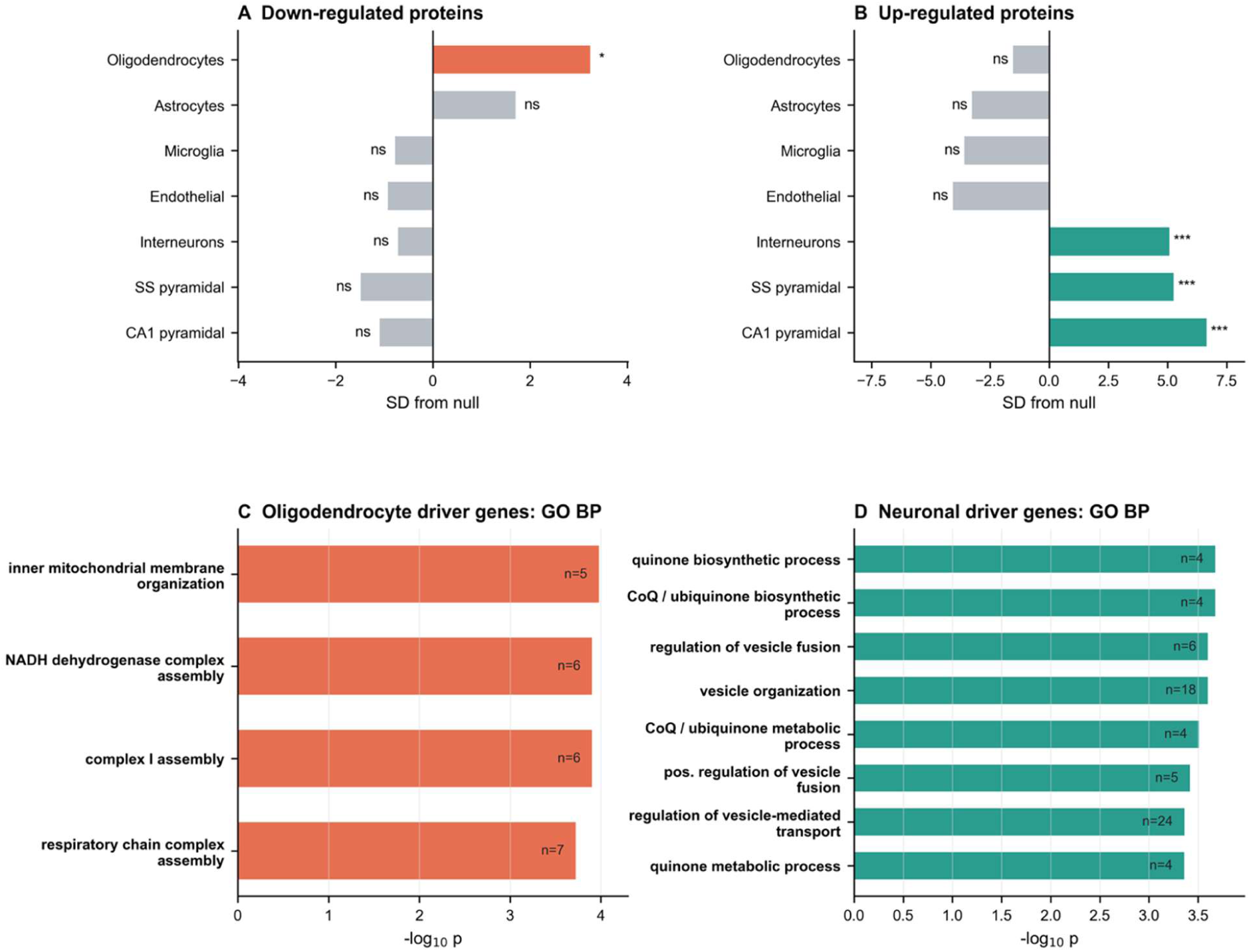
Cell-type enrichment of the LOU hippocampal signature and driver-gene pathway enrichment. A. EWCE of downregulated LOU proteins. B. EWCE of up-regulated LOU proteins. Both panels use the same vertical cell-type order: oligodendrocytes, astrocytes, microglia, endothelial cells, interneurons, SS pyramidal neurons and CA1 pyramidal neurons. Bars show standard deviations from the null specificity distribution; colored bars indicate q<0.05. C. GO Biological Process enrichment among oligodendrocyte driver genes from the down-regulated EWCE signal, highlighting mitochondrial inner-membrane organization and respiratory-chain complex I assembly. D. GO Biological Process enrichment among neuronal driver genes from the up-regulated EWCE signal, highlighting synaptic-vesicle and quinone / ubiquinone metabolism terms. Numbers inside bars indicate gene counts.

Pathway enrichment of the EWCE cell-specific driver gene lists confirmed compartment-specific function (Fig. 3C). Oligodendrocyte driver genes in the down-regulated set were dominated by complex I assembly and ETC components, suggesting that the OXPHOS-down signal of Section 3.2 originates in oligodendrocyte mitochondria. Importantly, bulk myelin proteins (MBP, PLP1, MAG, MOG, CNP, PMP22, PLLP, MOBP, CLDN11, ASPA, UGT8) were only modestly and non-significantly reduced in LOU (range −0.02 to −0.84), with a single MBP isoform reaching nominal significance (UniProt A0A8L2QCS8, log₂FC=−0.84, p=0.034).

Pathway enrichment on the neuronal driver genes in the up-regulated set (Fig. 3D) fell into two functional modules: synaptic, vesicular and membrane-binding terms, consistent with the SNARE and Hallmark Protein Secretion signals reported above; and quinone- and electron-carrier terms (FAD/NADH-related), consistent with sustained physiological electron flow at the neuronal cytoplasm/mitochondrion interface.

This oligodendrocyte phenotype indicates a quieter mitochondrial baseline in present, structurally intact oligodendrocytes, as opposed to one of cell loss or demyelination. This distinction will become important in the cross-species comparison (Section 3.7).

### 3.4 Synaptic vesicular fusion and inhibitory transmission elevated in LOU

At single-protein level, the neuronal up-regulation in LOU is divided in two thematically coherent groups. The first is the SNARE/vesicular fusion machinery: VAMP1 (log₂FC=+0.45, p=0.009), STX7 (+0.30, p=0.006), NSF (+0.21, p=0.012), VAMP7, NAPB, NAPG, with concurrent down-regulation of the ER–Golgi component STX18 (−0.24, p=0.046). These proteins constitute the core machinery of fast Ca2+-triggered exocytosis and vesicle recycling (Jahn et al., 2024; Sudhof & Rothman, 2009). The second is inhibitory transmission: the GABA_A receptor subunit GABRA3 was up-regulated (log₂FC=+0.51, p=0.037), consistent with a contribution to balanced excitatory–inhibitory tone (Olsen & Sieghart, 2009). Synaptic and vesicular machinery is among the most consistently reported molecular correlates of cognitive aging in humans, with progressive loss across the lifespan (Burke & Barnes, 2006; Fjell et al., 2014). This single-protein pattern matched the neuronal driver-gene enrichment for synaptic-vesicle, membrane-binding and electron-carrier terms identified by EWCE pathway analysis (Fig. 3D), supporting a neuronal program of elevated vesicular release and balanced excitatory-inhibitory signaling at baseline.

### 3.5 CD200 down with no inflammatory correlate

The strongest single-protein effect in the cleaned LOU set was a reduction in the neuronal CD200 ligand (log₂FC=−3.99, p=0.003), with no comparably reduced cell-surface neuronal protein elsewhere in the dataset. CD200 is broadly expressed on neurons and engages CD200R1 on microglia to restrain microglial activation (Hoek et al., 2000; Wright et al., 2003). Loss of this inhibitory signal in CD200-deficient mice or with age has been linked to microglial activation, reinforced inflammatory responses and increased vulnerability to insult (Cribbs et al., 2012; Lyons et al., 2007; Walker et al., 2009). In LOU, however, the CD200 reduction sits in a proteomic context with no upregulation of complement (C1q, C3, C4), no microglial activation marker (Aif1/Iba1, Trem2, Tyrobp), no astrogliosis (Gfap was at log₂FC=−0.52) and no inflammatory cytokine signature. In the global proteomic landscape, CD200 appears as an isolated extreme down-regulated protein rather than as part of a broader inflammatory shift (Fig. 1). This pattern is compatible with the absence of overt inflammatory activation at baseline, although microglial responsiveness was not directly assessed.(Norden & Godbout, 2013).

### 3.6 Hippocampal molecular divergence is more pronounced than cortical divergence in LOU rats

To understand whether the hippocampal signature reflects a brain-wide LOU phenotype or a regional one, we analyzed cortical proteome samples of another WIS-aCSF and LOU-aCSF cohort (Gephine et al., 2025) at the same age. The cortical landscape was substantially attenuated. Fewer proteins survived the same threshold, no cell-type signature reached significance by EWCE, and only a weak negative GSEA enrichment for respiratory chain remained (Fig. S2). The cortex-hippocampus intersection (Fig. S3) showed a small shared set, a much larger hippocampus-only set with neuronal EWCE enrichment and KEGG enrichment for energy and oxidative pathways, and a smaller cortex-only set without coherent cell-type assignment. At the age examined here, molecular divergence was substantially more pronounced in hippocampus than in cortex, suggesting that hippocampal pathways may be particularly sensitive to the biological processes distinguishing LOU and WIS rats. This observation is consistent with its known vulnerability in aging and AD (Kim et al., 2019; Wang et al., 2024) and with the cognitive readouts that dominate the LOU literature (Kollen et al., 2010).

### 3.7 Cross-species comparison: the LOU hippocampus is the directional inverse of late Alzheimer’s hippocampus

Because the LOU signature involved molecular systems strongly affected in AD, we investigated this axis by overlaying LOU log₂FCs onto our published 770-gene human hippocampal neuroinflammation panel applied to 30 human hippocampi spanning Braak 2 to Braak 6: 6 controls (Braak 2-3, CT), 12 early-AD (Braak 4, EAD) and 12 late-AD (Braak 5-6, LAD).

447 of the LOU-quantified proteins matched panel-curated genes; 162 had a published significant LAD-vs-CT log₂FC. Across these 162 genes, the LOU and LAD log₂FCs were anti-correlated (Spearman ρ=−0.157, p=0.046; Fig. 4). Directional testing supported this inverse relationship: 118 of 162 shared genes (72.8%) moved in opposite directions, while 44 of 162 (27.2%) moved in the same direction in LOU and late AD (binomial sign test p=5.3×10⁻⁹ against the 0.5 null). Restricting the analysis to LOU-significant genes (n=18), 14 of 18 (78%) still moved in the opposite direction, with 4 of 18 moving in the same direction (22%, two-sided exact binomial sign p = 0.031), supporting the inversion within the high-confidence subset. The LOU-vs-EAD comparison did not reach statistical significance at the whole-panel level (ρ=+0.20, p=0.23, n=37), reflecting both the smaller EAD DEG set and a different relationship between LOU and EAD on the oligodendrocyte axis.

**Figure 4.**
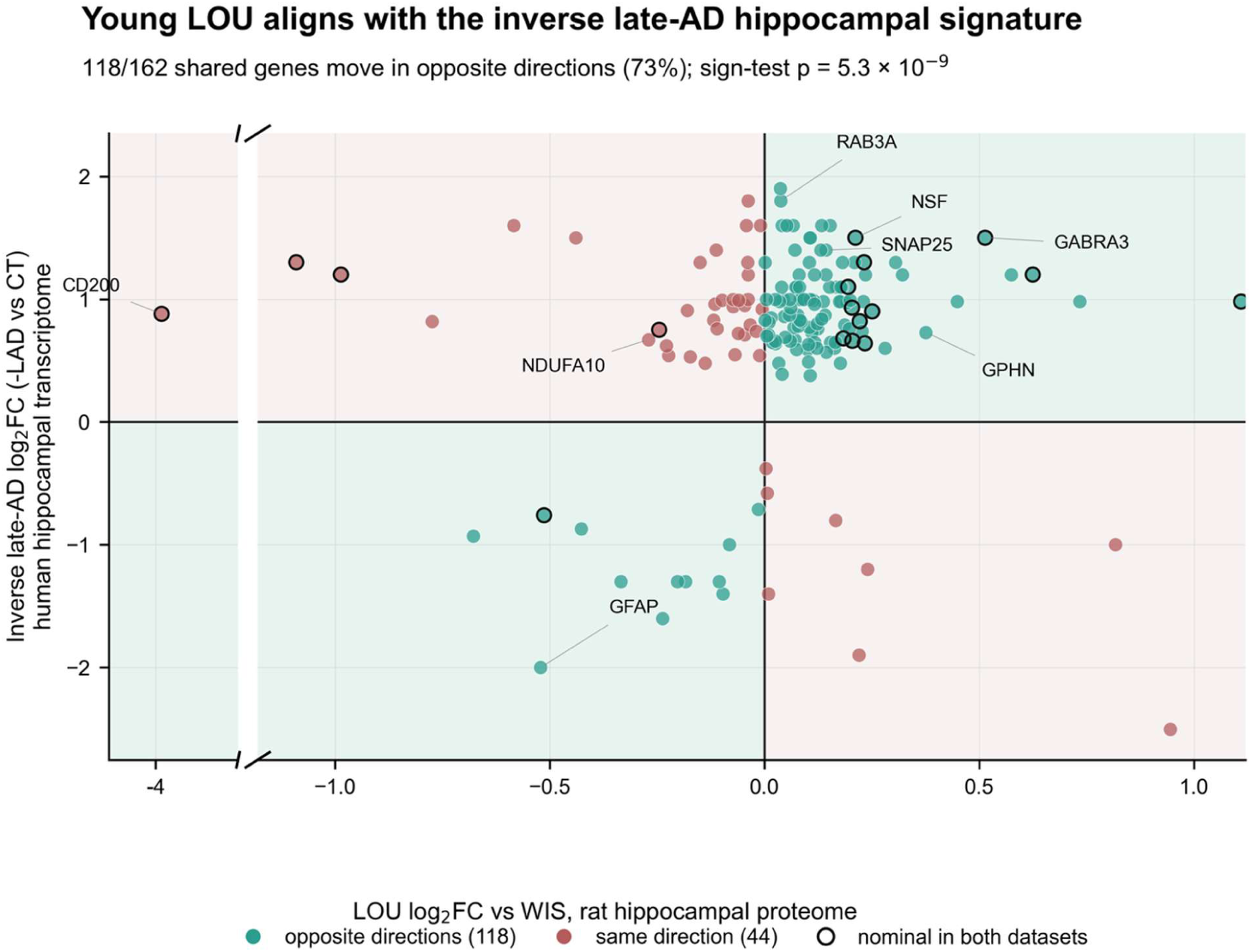
LOU aligns with the inverse late-AD hippocampal signature. LOU log₂FC (*vs* WIS; proteomics) is plotted against inverse LAD log₂FC (−LAD *vs* CT; human neuroinflammation transcriptomics) for the 162 shared genes. The same-sign quadrants after LAD inversion contain the 118 genes (72.8%) that move in opposite directions in the original LOU and LAD contrasts, whereas 44 genes (27.2%) move in the same direction. The sign test against the 0.5 null gives p=5.3×10⁻⁹; Spearman ρ=−0.16, p=0.046. Labelled genes highlight the opposing synaptic, mitochondrial, oligodendrocyte/myelin and neuro-immune components. The compressed x-axis preserves the full LOU range while exposing the dense central structure.

The neuro-immune axis is occupied at opposite poles in LOU and AD. Direct CD200 contrasts first show that CD200 decreases in LOU (log₂FC=−3.99, p=0.003) and in late AD (LAD *vs* CT log₂FC=−0.88, p=0.015; Fig. 5A). In LOU, GSVA scoring confirms that neuronal/synaptic, homeostatic microglial, DAM and central/peripheral immunity modules remain close to zero (Fig. 5B). In late AD, the same CD200 decrease occurs beside a broader neuronal/synaptic decrease (LAD−CT GSVA=−0.52, Tukey p = 0.031) and significant DAM and central/peripheral immunity increases (LAD−CT GSVA=+0.49 and +0.65; Tukey p = 0.024 and 0.009), against the previously reported complement up-regulation, astrogliosis, microglial activation and peripheral immune infiltration in our human cohort (Badina et al., 2025) (Fig. 5 C, E). The human ssGSEA sensitivity analysis gave the same qualitative neuronal/synaptic decrease and DAM increase (Fig. S4). The same CD200 loss therefore carries opposite biological meaning in the two contexts: in LOU it sits as an isolated, neuron-derived recalibration of immune tone against a quiet microglial background; in late AD it forms part of a degeneration-coupled loss of neuron-to-microglia inhibitory signaling, within an inflammatory and glial-remodeling context.

**Figure 5.**
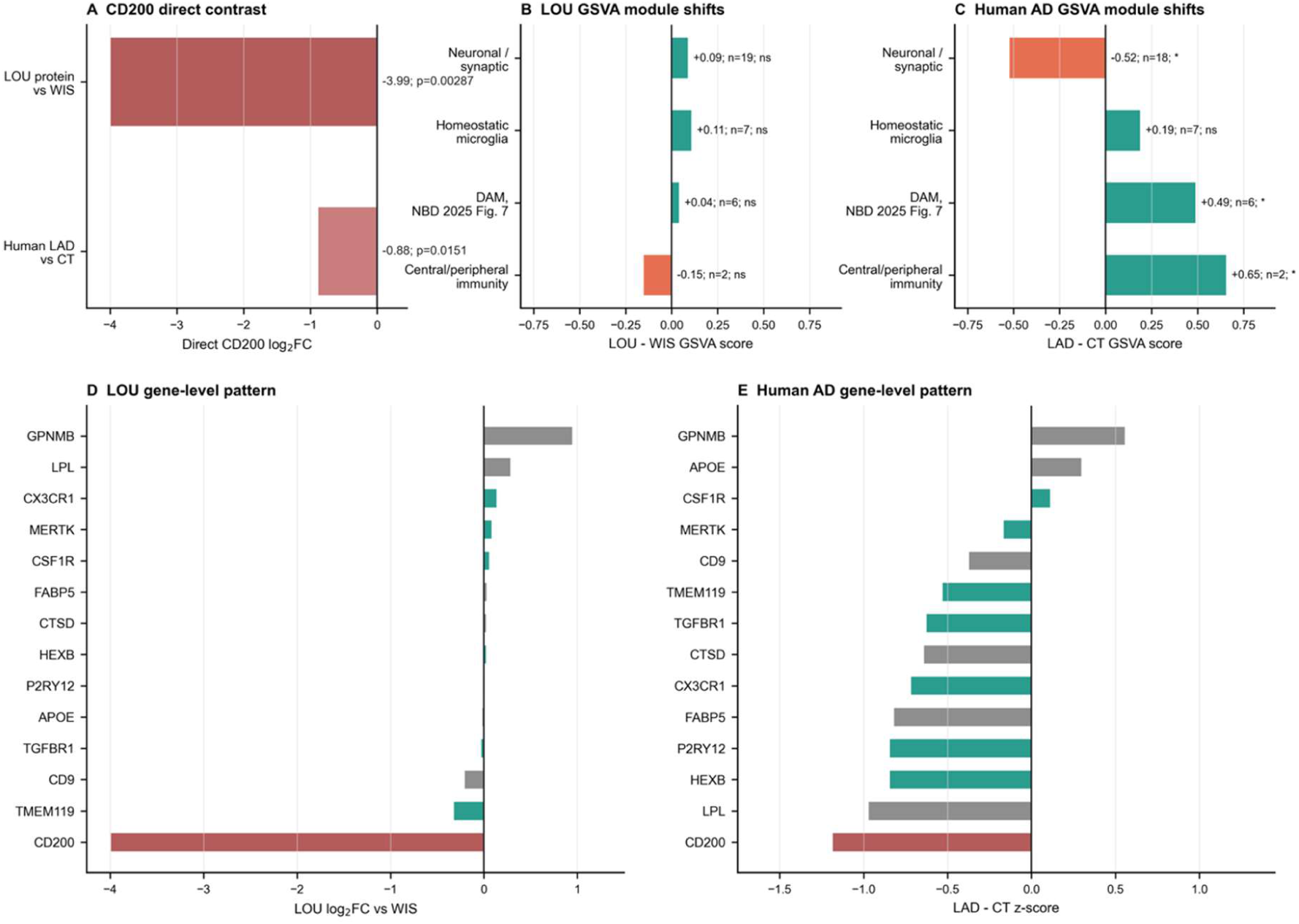
CD200 loss has opposite biological meaning in LOU and in human late AD. A. Direct CD200 contrast in the LOU vs WIS rat hippocampal proteome and the human LAD vs CT hippocampal transcriptome. B. GSVA module effects in LOU versus WIS rats, scored on the imputed hippocampal proteome after treating CD200 separately; neuronal/synaptic, homeostatic microglia, DAM and central/peripheral immunity modules remain close to zero. C. GSVA module effects in LAD versus CT human hippocampus, scored on the log₂ NanoString matrix; neuronal/synaptic scores decrease, whereas DAM and central/peripheral immunity scores increase. D. Gene-level LOU pattern for the shared CD200, homeostatic microglial and DAM panels, showing an isolated CD200 decrease. E. Gene-level LAD-CT z-score pattern for the same detected CD200, homeostatic microglial and DAM genes, showing that CD200 decrease occurs in an altered microglial context. Bars in A show direct log₂FC; bars in B-C show group-score differences; bars in D-E show gene-level contrasts. Labels in A give log₂FC and nominal p value; labels in B-C give effect size, number of detected/shared genes and significance (* p<0.05; ** p<0.01; *** p<0.001; ns p≥0.05). Gene sets were curated from the documented CD200 neuron-to-microglia axis, homeostatic/sensome microglial markers, the DAM panel from Badina et al. (2025), and central/peripheral immunity markers from the same human cohort.

The inversion was driven by neuronal categories in which LOU is up and late AD is down (Fig. 6). On the synaptic vesicular axis, four of four shared genes are up in LOU and down in late AD: NSF, SNAP25, RAB3A and SH3GL2, with LAD log₂FCs of −1.5, −1.4, −1.8 and −1.6. On the inhibitory synaptic axis, GABRA3 is the headline anti-concordant gene (LOU +0.51; LAD −1.50); the remaining inhibitory markers (GABRA1, GABRG2, GAD1, GAD2, GPHN, SHC3) cluster near zero in LOU but drop strongly in late AD, consistent with a higher LOU baseline that AD subsequently loses. On the general synaptic axis (NEFL, CAMK4, CNTNAP2, NMNAT2, HCN1, KCNAB1, PDE1A, DNM1, DNM3, ENO2), LOU effects are modest against strong LAD reductions, the same baseline-vs-collapse pattern. Side-by-side KEGG pathway enrichment supports the same conclusion. LOU-up KEGG terms (Aminoacyl-tRNA biosynthesis, Amino sugar metabolism and the V-ATPase term) match LAD-down KEGG terms in the same human cohort: GABAergic synapse (q=0.0014), Synaptic vesicle cycle (q=0.0068) and Oxidative Phosphorylation (q=0.0068, driven on the human side by ATP5/ATP6V/NDUFA10 transcripts and on the LOU side by the same V-ATPase, ARS and ETC machinery in the opposite direction (Table S5)).

**Figure 6.**
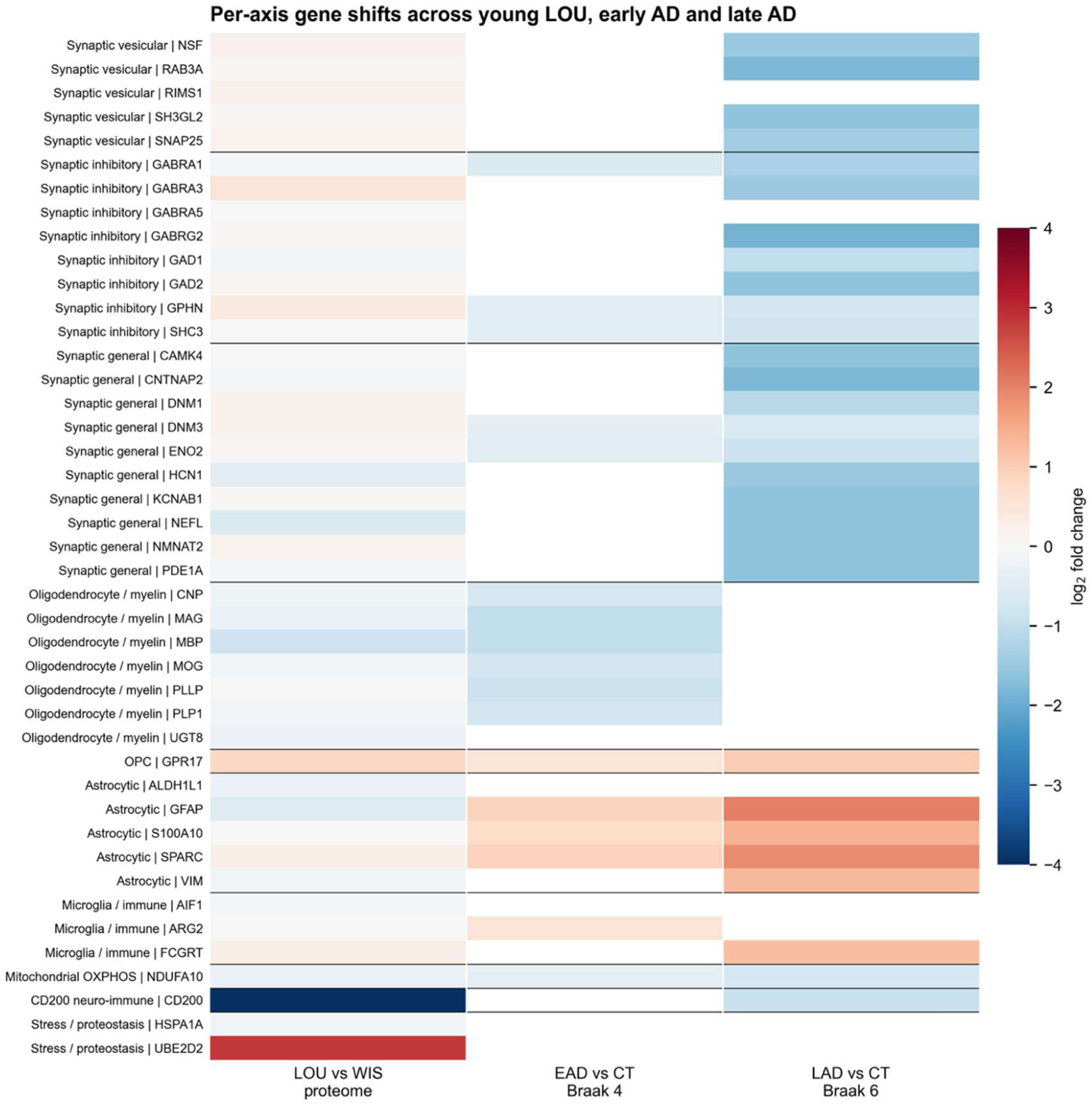
Per-axis log₂FC across LOU, early AD and late AD for genes present on both the human neuroinflammation panel and the LOU proteome. Columns show LOU vs WIS rat proteomics, EAD *vs* CT (Braak 4) and LAD vs CT (Braak 6) human transcriptomic contrasts. GFAP is assigned only to the astrocytic block. AIF1, ARG2 and FCGRT are retained in the microglia / immune block, reflecting microglial or Fc-immune signal rather than astrocytic identity. The heatmap shows the synaptic inversion, the oligodendrocyte / myelin continuum, the AD astrocytic response and the isolated CD200 decrease in LOU.

The oligodendrocyte and myelin axis behaved differently (Fig. 6, Oligodendrocyte / myelin block). Six of six shared myelin-associated genes (MAG, MBP, MOG, CNP, PLP1, PLLP) moved in the same direction in LOU and EAD (sign-test p=0.031). However, this directional overlap occurred in a distinct biological context: LOU showed no microglial activation, no complement up-regulation, no peripheral immune infiltration and no astrogliosis, whereas all four are features of EAD in our published neuroinflammation human cohort.

Each LOU axis found its opposite pattern in the human AD hippocampus. The mitochondrial axis sat on the same continuum as AD, with LOU at the quiescent end of OL OXPHOS modulation and AD at the failure end. The neuro-immune axis was occupied at opposite poles by LOU and AD. The synaptic axis was a directional inverse across LOU and late AD.

## 4. DISCUSSION

This study is the first comprehensive proteomic characterization of the hippocampus in adult LOU rats and the first direct comparison between this rodent model of successful brain aging and the human Alzheimer’s hippocampus. Three observations stand out at 3 months of age. Oligodendrocytes carry fewer nuclear-encoded complex I and III subunits while neurons carry more mitochondrial aminoacyl-tRNA synthetases and more lysosomal V-ATPase machinery. The neuronal CD200 ligand is strongly reduced against an otherwise non-activated glial and complement background. The SNARE machinery and the inhibitory GABA_A receptor subunit GABRA3 are coordinately elevated in pyramidal neurons and interneurons.

When mapped onto our published human hippocampal cohort spanning Braak 2 to Braak 6, this successful aging signature contrasts with the late-AD transcriptome along the synaptic, inhibitory neuron and neuro-immune communication axes. It is also associated with preserved oligodendrocyte and OXPHOS programs, whereas breakdown of these programs characterizes early AD.

### 4.1 Mitochondrial adaptations and oligodendrocyte energy metabolism

Our pathway and cell-type analyses seem to indicate a coordinated mitochondrial reconfiguration in LOU. Eleven nuclear-encoded complex I and III subunit genes form the leading edge of the GSEA Oxidative Phosphorylation result, with no mtDNA-encoded subunit included.

As EWCE places the downregulated OXPHOS signals in oligodendrocytes, these findings suggest a distinct mitochondrial configuration in oligodendrocytes, although the underlying structural organization of the respiratory chain cannot be determined from the present data. A reduced complex-I/OXPHOS protein footprint may be associated with altered electron transport chain dynamics and redox regulation. However, respiratory flux, electron leak and ROS production were not assessed in the present study (Barja, 2007; Caro et al., 2008; Lambert et al., 2010; Miwa et al., 2014; Sanz et al., 2005). This possibility is biologically plausible given the established contribution of complex I and complex III to mitochondrial superoxide production, but remains to be experimentally validated in the LOU hippocampus (Brand, 2016; Murphy, 2009), and with reports of reduced skeletal-muscle mitochondrial H₂O₂ production and oxidative-stress-related adaptations in the LOU strain (Garait et al., 2005; Piec et al., 2005). A possible functional consequence would be lower oxidative damage per unit ATP produced before aging has occurred, although this has not been measured directly in LOU hippocampal tissue. Supporting the possibility of altered redox regulation, the quinone-metabolism enzyme NQO2, which has been linked to ROS-related pathways, was among the strongest down-regulated proteins in the cleaned dataset. The functional significance of this decrease remains uncertain in the absence of direct oxidative stress measurements (Benoit et al., 2010; Gould et al., 2023; Janda et al., 2024; Janda et al., 2023).

These proteomic findings are consistent with previous transcriptomic data in the same strain. In a hippocampal microarray study, differential expression of genes involved in lipid metabolism, nuclear architecture, immune regulation and mitochondrial processes in old LOU rats compared to WIS was reported (Paban et al., 2013). The convergence between transcriptomic and proteomic datasets supports the view that mitochondrial metabolism and redox regulation are stable, multi-level hallmarks of the LOU phenotype.

### 4.2 Mitochondrial translation, lysosomal signaling and mTORC1

In parallel to the oligodendrocyte OXPHOS-down signal, 27 of 42 leading-edge aminoacyl-tRNA synthetase genes are mitochondrial, and the V-ATPase, LAMTOR and mTORC1 machinery is up-regulated. EWCE places the V-ATPase, SNARE and electron-carrier signal in pyramidal neurons and interneurons.

Accurate translation of the few mtDNA-encoded subunits remaining on each chain (Couvillion et al., 2016; Sissler et al., 2017), and matched anabolic and vesicular machinery downstream (Saxton & Sabatini, 2017; Zoncu et al., 2011), would be required if flux is maintained despite a smaller respiratory-chain protein footprint. Aminoacyl-tRNA synthetase deficits cause neurometabolic disorders and impaired energy homeostasis (Fine et al., 2019; Gonzalez-Serrano et al., 2019); Therefore, their up-regulation in LOU rats may reflect an increased investment in translational machinery. Whether this translates into improved translation fidelity, proteostasis or synaptic function remains to be determined. The up-regulation of several V-ATPase subunits (ATP6V1G2, ATP6V1C1, ATP6V1E1) together with LAMTOR3 suggests potential alterations in lysosomal and nutrient-sensing pathways. However, neither lysosomal activity nor mTORC1 signaling were directly assessed in this study.(Saxton & Sabatini, 2017; Zoncu et al., 2011). V-ATPase up-regulation is, however, also compatible with autophagic flux, and the two programs are partly competing. The concurrent up-regulation of mitochondrial aminoacyl-tRNA synthetases, SNARE-associated proteins and MSigDB Hallmark Protein Secretion suggests increased biosynthetic activity. However, the relative balance between anabolic and degradative processes cannot be established from proteomic data alone.

### 4.3 Synaptic signaling, vesicular dynamics and inhibitory tone

Beyond mitochondrial metabolism, the LOU hippocampus exhibits significant modulation of SNARE complex components involved in synaptic vesicle fusion and recycling. Increased levels of VAMP1, NSF and the SNAP family proteins indicate remodeling of presynaptic molecular machinery. Whether these changes translate into altered neurotransmitter release, vesicle recycling efficiency or synaptic plasticity remains unknown (Jahn et al., 2024; Sudhof & Rothman, 2009). The upregulation of GABRA3 suggests modifications of inhibitory signaling pathways, although the consequences for network activity and excitation–inhibition balance were not directly examined. The decrease in Syntaxin-18, a component of the ER-Golgi trafficking pathway, may reflect optimized protein folding and reduced ER stress, processes often disrupted during aging. Together, these changes identify a coordinated synaptic molecular signature involving vesicle trafficking and inhibitory signaling pathways, although the functional consequences remain to be established.4.4 A primed neuro-immune set-point

The strongest single-protein effect in LOU is the reduction of the neuronal CD200 ligand (log₂FC=−3.99, p=0.003). CD200 is constitutively expressed on neurons and engages microglial CD200R1 to maintain microglia in a quiescent surveillance state (Hoek et al., 2000; Wright et al., 2003); reduction of CD200 with age has been used to explain microglial priming and the heightened inflammatory response of the aged hippocampus (Cribbs et al., 2012; Lyons et al., 2007; Walker et al., 2009).

In LOU, the CD200 reduction sits in a proteomic context with no complement up-regulation, no astrogliosis, no microglial activation marker and no inflammatory cytokine signature; GSVA makes the same point at module level, with flat neuronal/synaptic, homeostatic microglia, DAM and central/peripheral immunity scores alongside the CD200 decrease. In LAD, the same CD200 decrease appears in the opposite configuration, with neuronal/synaptic loss and DAM/immunity enrichment. One possible interpretation is that reduced CD200 expression reflects an altered neuroimmune homeostatic state. However, the functional status of microglia cannot be inferred directly from the present data.(Norden & Godbout, 2013).

### 4.5 Hippocampal specificity compared with cortex

The hippocampal signature does not appear to reflect a uniform brain-wide LOU phenotype at 3 months of age. In the cortical proteome, the landscape was substantially attenuated. Fewer proteins survived the same threshold, no cell-type signature reached significance by EWCE, and only a weak negative GSEA enrichment for respiratory chain remained. The cortex-hippocampus showed a small shared set, a much larger hippocampus-only set with neuronal EWCE enrichment and KEGG enrichment for energy and oxidative pathways, and a smaller cortex-only set without coherent cell-type assignment.

The hippocampus thus appears as the site of earliest LOU divergence at 3 months of age, consistent with its known vulnerability in aging and AD, and with the cognitive readouts that dominate the LOU literature. This regional specificity also supports the relevance of comparing the LOU hippocampal proteome with the human Alzheimer’s hippocampus.

### 4.6 Super-aging and pathological aging share an axis structure

When projected onto the human Alzheimer’s hippocampus, this mitochondrial and oligodendrocyte configuration aligns with the same broad molecular axis affected in pathological aging, but in a different biological state. Six shared myelin-associated genes (MAG, MBP, MOG, CNP, PLP1, PLLP) move in the same direction in LOU and early AD, but in LOU this occurs without microglial activation, complement up-regulation, or astrogliosis. KEGG Oxidative Phosphorylation moves in opposite directions across the two species, driven on the LOU side by oligodendrocyte down-regulation of nuclear-encoded complex I/III subunits, and on the human side by ATP5/ATP6V/NDUFA10 transcripts in late AD. LOU oligodendrocytes therefore exhibit a molecular profile compatible with reduced inflammatory engagement and altered mitochondrial regulation relative to the AD-associated signature. Whether this reflects a lower cellular stress state remains to be determined experimentally (Badina et al., 2025).

The same synaptic configuration is the strongest single contributor to the cross-species inversion. In late AD, NSF, SNAP25, RAB3A, SH3GL2, GABRA3 and the broader GABAergic synapse and synaptic vesicle cycle programs all drop sharply; in LOU these same proteins and pathways are elevated at baseline. The 5.3×10⁻⁹ sign test against the 0.5 null over 162 shared genes (Fig. 4) confirms that the resilience phenotype acts on the same molecular machinery that pathology disturbs, in the opposite direction.

The cross-species comparison narrows the interpretation of the neuro-immune axis. In EAD, microglia are at the activated end of the same axis, with strong microglial and complement up-regulation (Badina et al., 2025); in LOU CD200 is reduced without a parallel inflammatory program. Module-level analysis (Fig. 5) makes this contrast explicit: in LOU, CD200 is the only strongly shifted node, while homeostatic microglia, DAM and neuronal/synaptic modules are flat; in late AD, the same CD200 reduction sits inside a configuration of falling neuron / synapse markers and DAM/immunity enrichment. CD200 down-regulation thus carries opposite biological meaning across the two contexts: an isolated recalibration of immune tone in LOU, versus a component of degeneration-coupled loss of neuron-to-microglia inhibitory signaling in AD. Taken together, the four axes show that super-aging in the rat and pathological aging in humans act on the same molecular axes, in opposite directions. The synaptic and inhibitory axis is a strict directional inverse. The neuro-immune axis is occupied at opposite poles, with LOU lowering CD200 in a quiet glial environment and AD up-regulating microglial and complement effectors in an inflammatory one. The oligodendrocyte axis defines a continuum on which LOU sits at the quiescent end and EAD at the failure end; LAD partially recovers myelin transcripts as inflammation subsides (Badina et al., 2025), matching the recovered, low-stress oligodendrocyte state that LOU appears to occupy throughout. The phenotypes are not separate. They are opposite ends of the same hippocampal molecular spectrum. These findings identify a hippocampal molecular profile associated with successful aging that shows broad inverse relationships with molecular signatures reported in Alzheimer’s disease. Whether these opposing patterns reflect distinct positions along a shared biological continuum remains an open question.

### 4.7 Limits and perspectives

Several limitations should be considered. First, the hippocampal proteomic comparison was performed in a small cohort of male rats (n=4 per group), and no individual protein survived Benjamini–Hochberg correction at q<0.10. Accordingly, single-protein changes should be interpreted as nominal observations. The main inference therefore relies on convergent evidence across three complementary analytical levels: pathway-level GSEA, EWCE cell-type enrichment and cross-species directional analyses.

Second, the study captures an adult pre-aging time point. It identifies a baseline molecular configuration associated with the LOU strain, but does not determine whether this configuration is maintained, amplified or remodeled during aging. Longitudinal profiling at 6, 12, 18 and 24 months in the same strain pair would be required to test whether the LOU molecular signature remains stable, progressively strengthens, or drifts with age. Such analyses would also determine whether WIS oligodendrocyte-related changes progressively converge toward an early Alzheimer’s disease-like state.

Third, the cross-species comparison integrates rat hippocampal proteomics with a targeted human transcriptomic panel restricted to neuroinflammatory markers. It should therefore be interpreted as a directional overlay of shared molecular axes rather than as a direct quantitative equivalence between species or omic layers. For this reason, direction- and rank-based statistics were favored over direct effect-size matching across modalities. Whole-transcriptome RNA sequencing in the same human cohort, combined with broader proteomic profiling in rats, would strengthen the cross-species interpretation.

Fourth, the functional consequences of the mitochondrial, lysosomal, synaptic and CD200-related changes identified here were not directly tested. In particular, targeted CD200/CD200R1 perturbation in the LOU strain, combined with longitudinal cognitive assessment and multiplexed neuroinflammatory profiling, would help determine whether the CD200 down-shift contributes causally to a quiescent microglial set-point or merely accompanies it. Finally, the present study used only male rats, whereas the human cohort was mixed-sex. Whether the LOU resilience phenotype is sex-modified remains an open question.

By identifying early-life proteostatic, mitochondrial, synaptic and neuroimmune configurations that precede cognitive decline, the present study positions the LOU rat as a distinctive model of spontaneous, non-pathological cognitive longevity. It provides a tractable molecular framework for investigating the determinants of cognitive aging resilience. The cross-species comparison suggests that this resilience phenotype involves molecular axes that are disrupted in the human Alzheimer’s hippocampus, but in opposite directions. In this comparison, NDUFA9, UQCRC2, NQO2, NSF, GABRA3, the V-ATPase complement and CD200 emerge as candidate molecular nodes distinguishing resilient from pathological aging. Together, these findings support the hypothesis that successful cognitive aging and Alzheimer’s disease may reflect divergent configurations of partly shared molecular systems, raising the central mechanistic question of what drives the trajectory split between resilience and pathology.

## CREDIT AUTHOR CONTRIBUTIONS

Lucas Gephine: conceptualization, data curation, formal analysis, investigation, writing - original draft, writing - review & editing, visualization. Aurélien M. Badina: conceptualization, data curation, formal analysis, investigation cross-species comparative analysis, writing - review & editing, visualization. Sophie Corvaisier: conceptualization, data curation, formal analysis, writing - review & editing. Benjamin B. Tournier: conceptualization, supervision, validation, writing - review & editing. Marianne Léger: conceptualization, funding acquisition, project administration, supervision, visualization, writing - review & editing. Thomas Freret: conceptualization, funding acquisition, project administration, resources, supervision, validation, visualization, writing - review & editing.

## DECLARATION OF GENERATIVE AI USE

During the preparation of this work the authors used a large-language-model assistant to help organize analytical scripts (Python and R) and to review grammar and readability for selected sections of the Methods and Results. All analytical code was reviewed and tested by the authors against the source data; all numerical values reported were independently verified against the supplementary tables; all text was reviewed and revised by the authors, who take full responsibility for the content of the publication.

## DATA AVAILABILITY

Raw mass-spectrometry data and processed protein abundance tables are available from the corresponding authors upon reasonable request and will be deposited to the ProteomeXchange consortium via PRIDE before publication. Annotated DEP tables, enrichment outputs, cross-species overlay tables and analysis scripts are provided as Supplementary Material 2. The human hippocampal neuroinflammation cohort and its DEG tables were previously published (Badina et al., 2025); the relevant DEG outputs are reproduced for clarity in Supplementary Material 2 with explicit reference to the original source.

## STUDY APPROVAL

All animal procedures were approved by the Normandy Ethics Committee for Animal Experimentation (CENOMEXA) and conducted in compliance with EU Directive 2010/63 and the ARRIVE guidelines (APAFIS#38885-2022101410092500-v1). The human hippocampal cohort was obtained from the Netherlands Brain Bank under a previously approved protocol (Cantonal Commission for Research Ethics, Geneva; see Badina et al., 2025).

## ACKNOWLEDGMENTS

We thank the Proteogen Platform (SFR ICORE 4206, Unicaen, 14000 Caen, France) for proteomic data acquisition, and the Netherlands Brain Bank (Amsterdam, Netherlands) for human hippocampal tissue.

## SUPPLEMENTARY DATA

Table S1. Annotated full proteome (lou_hipp_all_proteins_annotated.csv): UniProt, GeneSymbol, P-value, log₂FC, BH q-value, contaminant flags, mt-Nd2 outlier flag, classification.

Table S2. Cleaned DEP set with directional split (lou_hipp_DEPs_clean_nominal.csv).

Table S3. Full ORA outputs by library × direction (enrichr/*).

Table S4. GSEA preranked outputs by library (gsea_prerank/*).

Table S5. Cross-species panel overlay table per gene (lou_panel_overlay_full.csv) and Spearman / sign-test summaries (lou_vs_ad_spearman.csv, lou_vs_ad_sign_test.csv).

Figure S1. EWCE driver-gene pathway enrichment (oligodendrocyte and neuronal drivers).

Figure S2. Cortical proteomic overview at 3 months (LOU-aCSF vs WIS-aCSF arms of RESILOU_STZ).

Figure S3. Cortex-hippocampus intersection (Venn diagram, region-only EWCE and KEGG). Figure S4. Sensitivity quantification of the human GSVA module effects by per-subject ssGSEA / GSVA-equivalent scoring.

