## Supplementary material for "Proteomic signatures of cognitive resilience in LOU/c/Jall rats converge with inverse hippocampal axes of Alzheimer’s disease"

*Successful aging and Alzheimer's disease: opposite ends of the same hippocampal molecular spectrum*

This supplementary material is organised into three sections:

• Supplementary Methods notes (cross-species directional concordance; DAM module methodology).

• Supplementary Figures S1-S4.

• Supplementary Tables S1-S5. Large tables are provided as companion .xlsx files referenced in the table headers below.

#### Supplementary Methods notes

##### S-Methods 1. Cross-species directional concordance: LOU-significant subset

In addition to the whole-panel concordance test reported in §3.7 of the main text (n = 162 shared genes, 118 opposite / 44 same direction, sign-test $p = 5.3 \times 10^{-9}$), we computed a sensitivity analysis restricted to LOU-significant genes. Specifically, of the 162 panel-overlapping genes with both a LOU $log_2FC$ and a published Braak 6 (LAD) − CT $log_2FC$, we retained those reaching nominal LOU significance (limma moderated p-value < 0.05). This subset comprised n = 18 genes; the binomial sign test against the 0.5 null was applied identically to the whole-panel analysis (two-sided). 4 of 18 (22 %) of these LOU-significant genes moved in the same direction in LOU and late AD (sign-test p = 0.031), confirming the inversion within the high-confidence subset.

##### S-Methods 2. CD200 / microglia module quantification: GSVA and ssGSEA sensitivity

Two related quantifications of the neuronal/synaptic, homeostatic-microglia, disease-associated microglia (DAM, Badina et al., 2025) and central/peripheral immunity modules are presented in this manuscript and its supplement. CD200 is shown separately as a direct single-gene/protein contrast in main-text Figure 5A.

(i) Exact GSVA (main-text Figure 5B-C). For LOU, module scores were computed on the $log_2$ imputed hippocampal protein matrix for WIS and LOU rats. For human AD, the same module scores were computed on the published $log_2$ NanoString expression matrix. Group contrasts are shown as LOU − WIS and LAD − CT score differences.

(ii) Per-subject ssGSEA / GSVA-equivalent (Supplementary Figure S4). For each human subject and each multi-gene module, we computed a single-sample gene-set enrichment score on the $log_2$-transformed NanoString expression matrix (757 endogenous probes × 30 subjects) using gseapy.ssgsea with rank-based sample normalisation. Group differences were assessed with a one-way ANOVA followed by Tukey's HSD test.

Exact GSVA is used for main-text Figure 5B-C because it captures per-subject module enrichment directly and recovers the DAM increase in LAD reported in Badina et al. 2025 Fig. 7D (Tukey CT-LAD p = 0.024). The ssGSEA sensitivity analysis in Supplementary Figure S4 gives the same directional interpretation for the human multi-gene modules. The underlying GSVA and ssGSEA outputs are deposited in cd200_microglia/gsva_analysis/.

### Supplementary Figures

##### Figure S1. EWCE driver-gene pathway enrichment

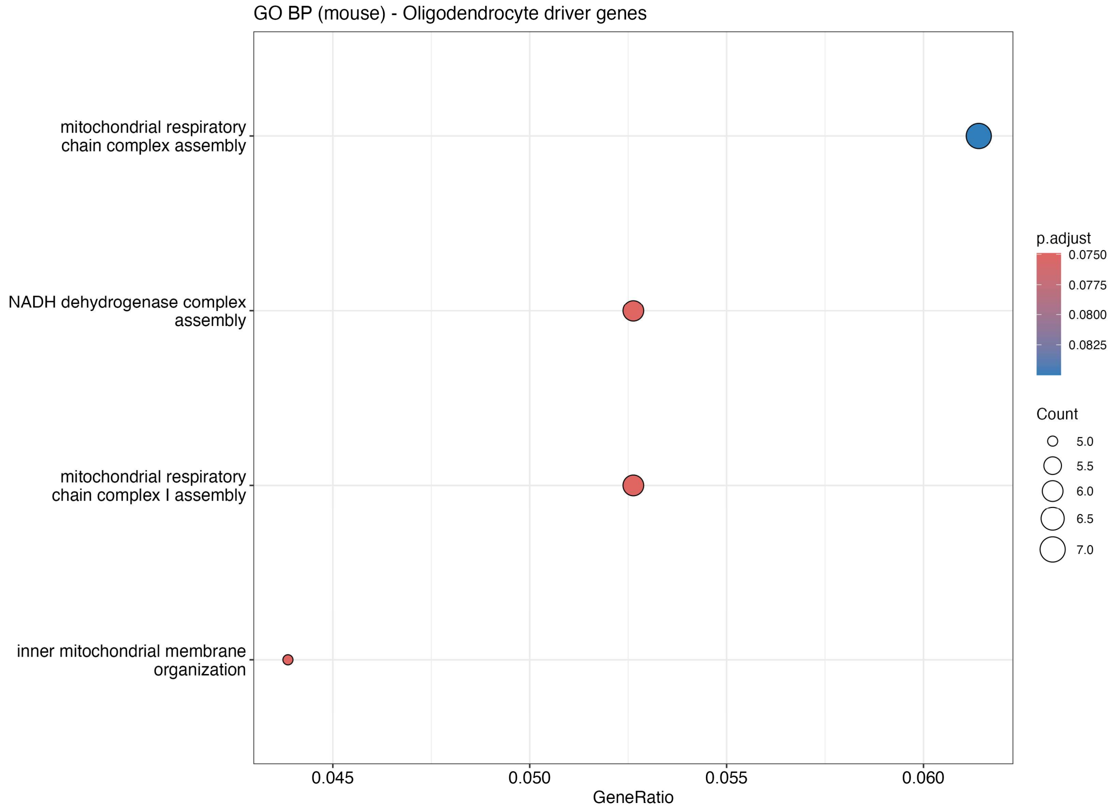

**Figure S1A.** GO Biological Process enrichment among the oligodendrocyte driver genes from the down-regulated EWCE signal. The dominant terms are mitochondrial inner-membrane organisation and respiratory-chain complex I assembly, indicating that the OXPHOS-down signal originates in oligodendrocyte mitochondria.

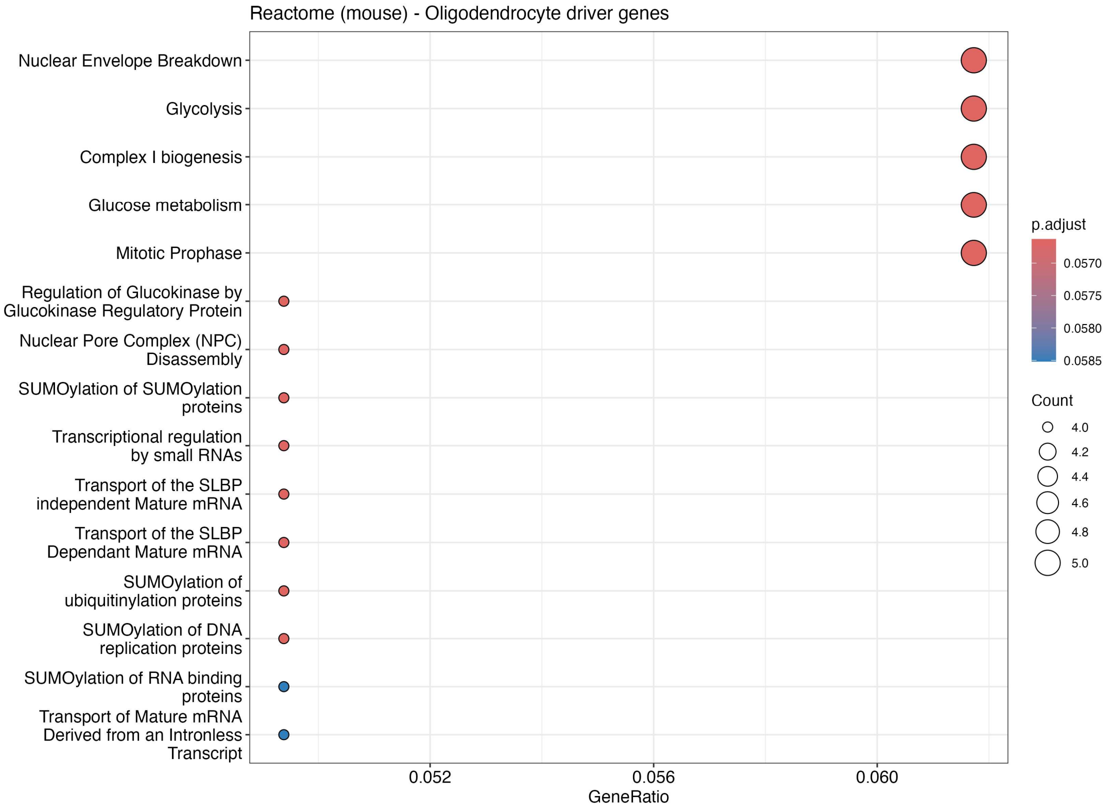

**Figure S1B.** Reactome enrichment among the oligodendrocyte driver genes from the down-regulated EWCE signal, confirming complex I assembly and electron-transport-chain components as the dominant Reactome pathways.

**Figure S2. Cortical proteomic overview at 3 months**

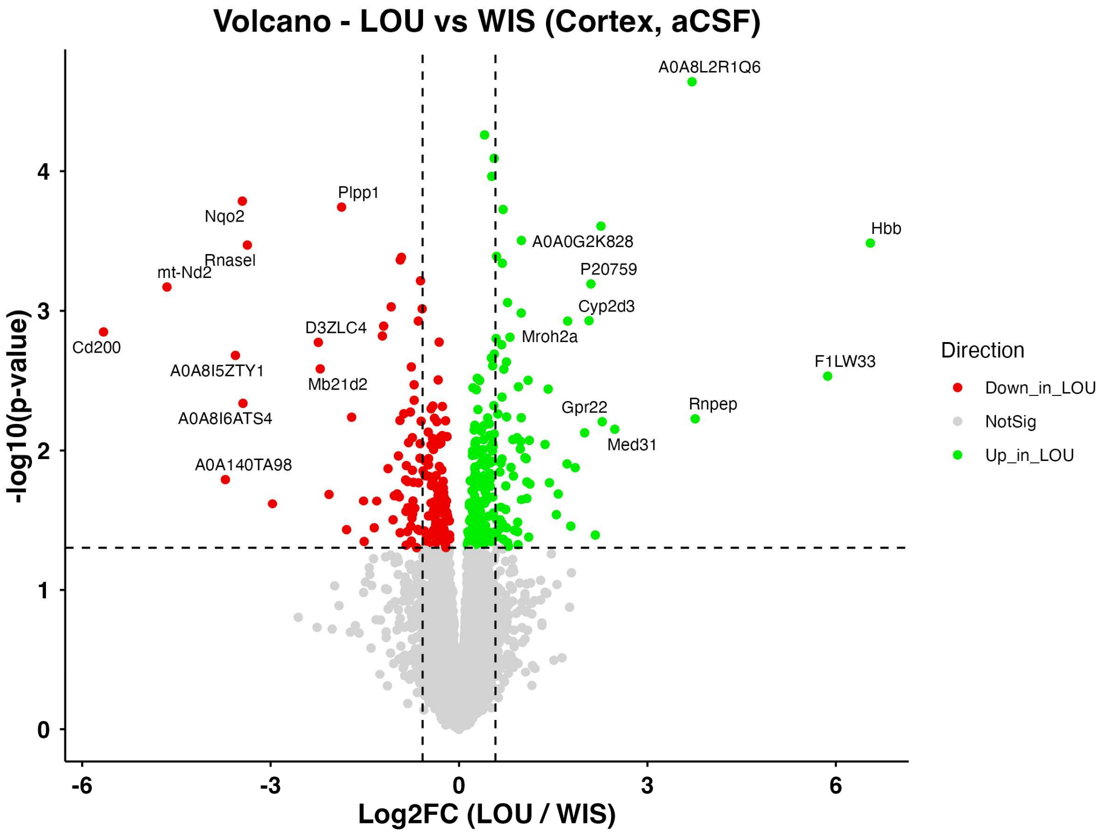

**Figure S2A.** Volcano plot of cortical LOU vs WIS proteins (WIS-aCSF and LOU-aCSF arms of RESILOU_STZ, 3 months, n = 7 per group). The cortical landscape is substantially attenuated compared with the hippocampal landscape (main-text Fig. 1).

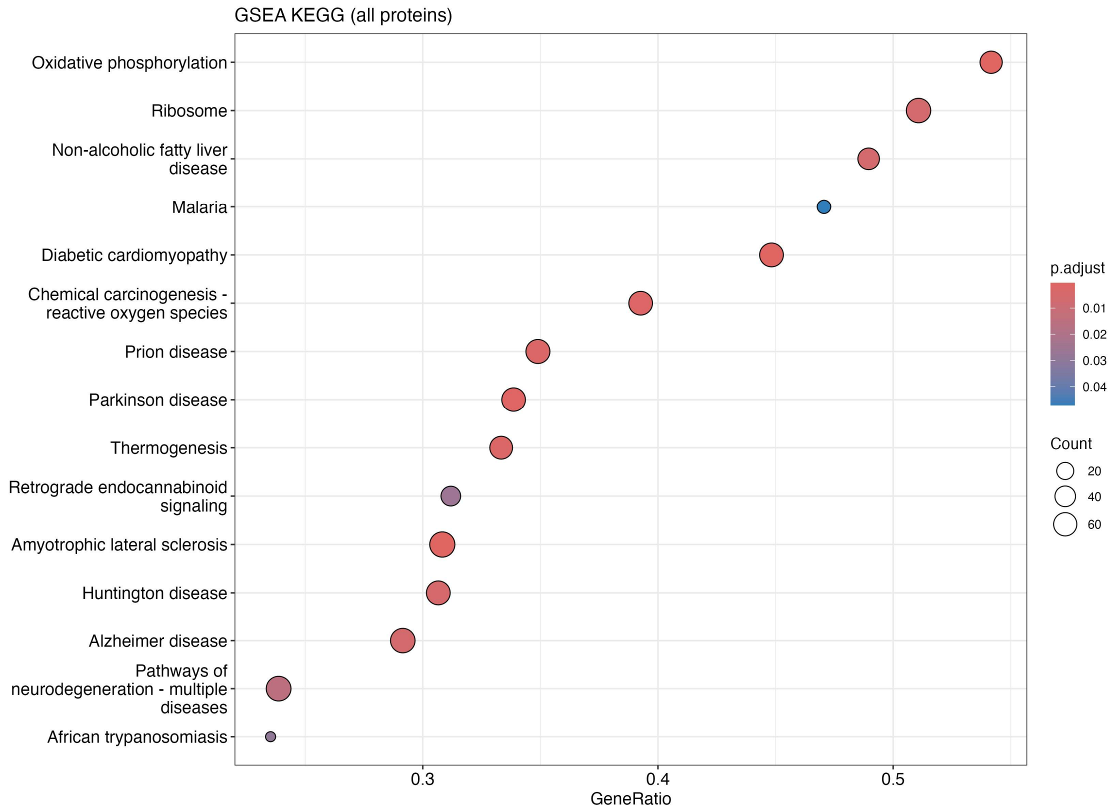

**Figure S2B.** KEGG pre-ranked GSEA on the cortical ranked proteome. Only a weak negative enrichment for respiratory-chain related sets remains.

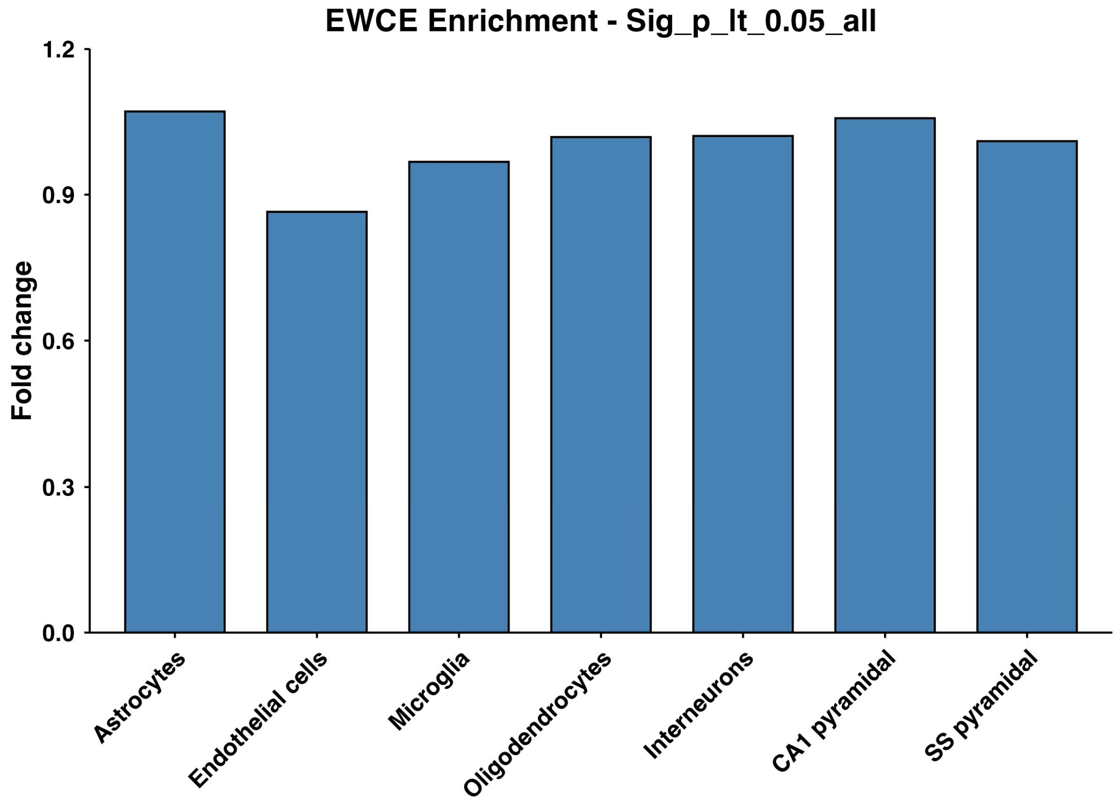

**Figure S2C.** EWCE of cortical DEPs against the ewceData mouse hippocampal single-cell reference at annotation level 1. No cell type reaches the q < 0.05 threshold in either direction.

**Figure S3. Cortex / hippocampus intersection**

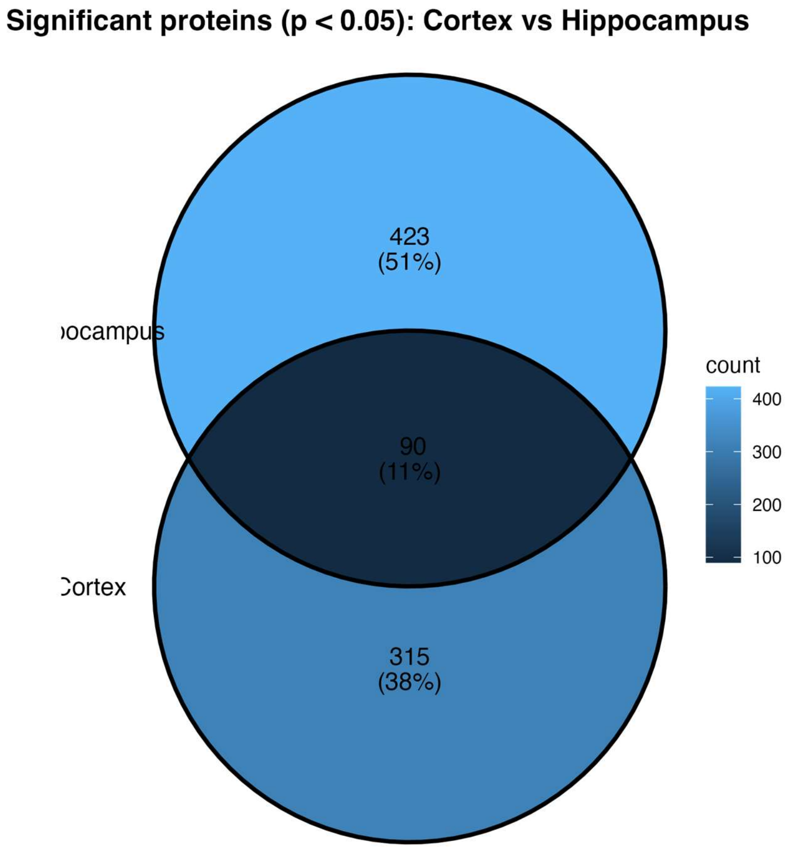

**Figure S3A.** Venn diagram of significant proteins ($p < 0.05$, $|\log_2 FC| \geq 0.18$) shared between the cortical (LOU-aCSF vs WIS-aCSF) and hippocampal (LOU vs WIS) DEP sets at 3 months.

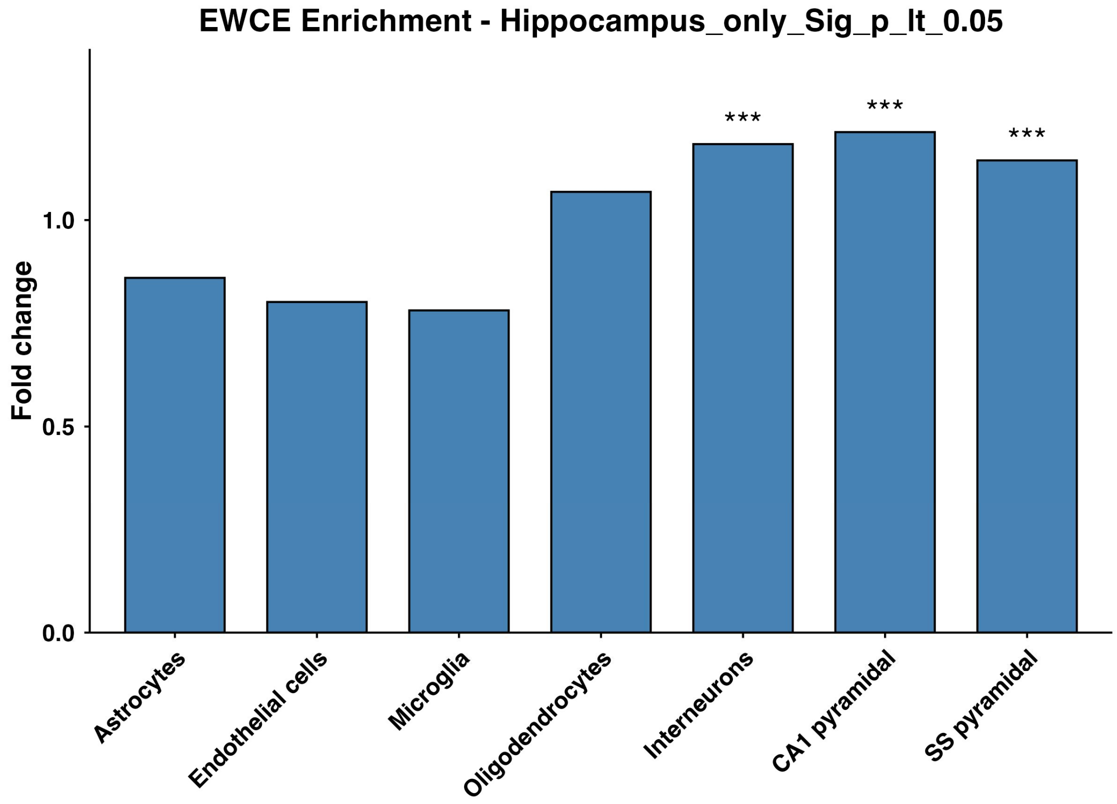

**Figure S3B.** EWCE on hippocampus-only DEPs. Pyramidal neuron and interneuron enrichments remain when cortex-shared proteins are removed.

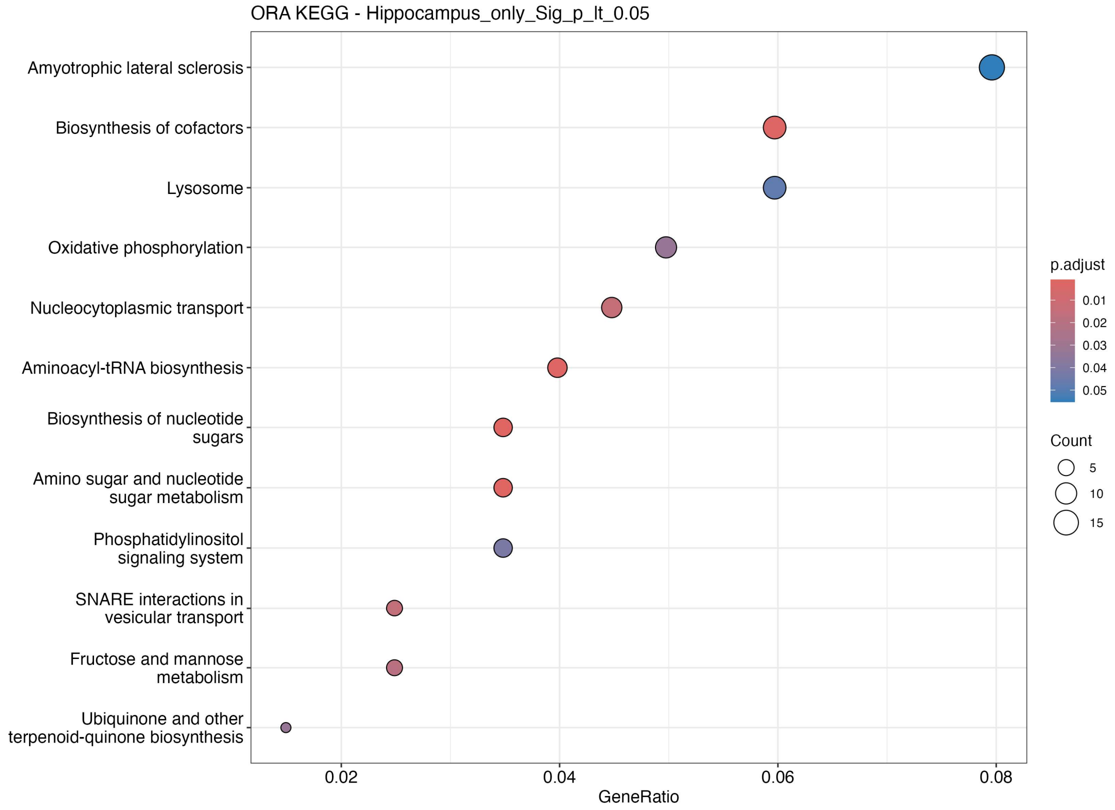

**Figure S3C.** KEGG enrichment on hippocampus-only DEPs. Aminoacyl-tRNA biosynthesis, V-ATPase and OXPHOS terms are retained.

#### Figure S4. Sensitivity quantification of Fig. 5C: per-subject ssGSEA / GSVA-equivalent

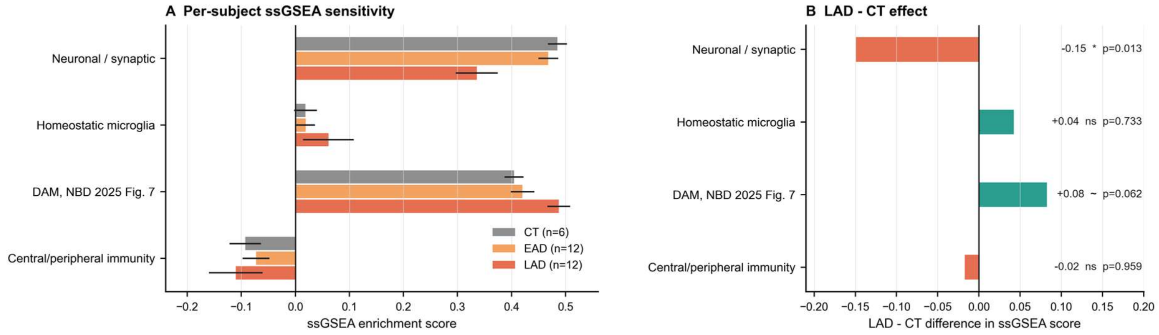

Figure S4. Per-subject ssGSEA / GSVA-equivalent scoring of the four human multi-gene modules shown in main-text Figure 5C (neuronal/synaptic, homeostatic microglia, DAM, central/peripheral immunity) on the $log_2$-transformed NanoString expression matrix (757 endogenous probes × 30 subjects). A. Mean enrichment score per group (CT, n = 6; EAD, n = 12; LAD, n = 12; bars are standard errors of the mean). B. LAD − CT effect with Tukey HSD significance markers (* $p < 0.05$; ** $p < 0.01$; *** $p < 0.001$; ~ $p < 0.1$; ns $p \geq 0.05$). The DAM module is directionally higher in LAD by global ANOVA (ANOVA p = 0.031; Tukey CT-LAD p = 0.062; Tukey EAD-LAD p = 0.065), matching the direction reported in Badina et al. 2025 Fig. 7D; the neuronal/synaptic module decreases significantly (ANOVA p = 0.003, Tukey CT-LAD p = 0.013, Tukey EAD-

LAD p = 0.0065). See S-Methods 2 for the relationship between this sensitivity analysis and the exact-GSVA main-text Figure 5C.

### Supplementary Tables

All tables are also provided as standalone .xlsx files in the submission_package/tables folder. Below, the metadata and a short preview are reproduced inline; full content lives in the companion spreadsheet for ease of programmatic access.

##### Table S1. Annotated full proteome

Source file: tables/Table_S1_annotated_proteome.xlsx (3 sheets: all_proteins n = 7 984; cleaned_DEPs n = 473; excluded_contaminants n = 35).

Columns: UNIPROT, P_value, log2FC, GeneSymbol, Entrez, Sig, Direction, negLog10P, q_value_BH, negLog10q, is_globin, is_ig, is_plasma, contaminant, uniprot_known_blood_ig, mt_nd2_outlier, set_class.

| UNIPROT | GeneSymbol | log2FC | P_value | q_value_BH | Direction | set_class |
|---|---|---|---|---|---|---|
| **A0A0G2JTJ9** | Tfcp2 | 0.533 | 6.03e-05 | 0.481 | Up_in_LOU | Sig_p |
| **D4ACS3** | Mb21d2 | -2.73 | 0.000434 | 0.524 | Down_in_LOU | Sig_p |
| **Q5XIB4** | Ufsp2 | 0.703 | 0.000528 | 0.524 | Up_in_LOU | Sig_p |
| **F7FBS0** | Bphl | 0.673 | 0.000532 | 0.524 | Up_in_LOU | Sig_p |
| **A0A8I6GJU0** | Rnpep | 2.72 | 0.000637 | 0.524 | Up_in_LOU | Sig_p |
| **A0A8I5Y874** | Adck1 | -0.934 | 0.000799 | 0.524 | Down_in_LOU | Sig_p |
| **A0A8I6A660** | Esyt2 | 0.429 | 0.00105 | 0.524 | Up_in_LOU | Sig_p |
| **Q64380** | Sardh | -0.896 | 0.00106 | 0.524 | Down_in_LOU | Sig_p |
| **A0A8I6AVP7** | Odr4 | 0.714 | 0.00112 | 0.524 | Up_in_LOU | Sig_p |
| **Q9R1B1** | Timm10b | -0.562 | 0.00121 | 0.524 | Down_in_LOU | Sig_p |
| **Q66HC3** | RGD1359108 | 0.284 | 0.00125 | 0.524 | Up_in_LOU | Sig_p |
| **Q8VHE9** | Retsat | -1.02 | 0.00126 | 0.524 | Down_in_LOU | Sig_p |
| **Q9Z339** | Gsto1 | 0.468 | 0.00145 | 0.524 | Up_in_LOU | Sig_p |
| **Q4V7A0** | Skic8 | 0.27 | 0.00154 | 0.524 | Up_in_LOU | Sig_p |
| **A0A0G2JSR2** | Syt3 | 0.659 | 0.00167 | 0.524 | Up_in_LOU | Sig_p |

*Top 15 of 473 cleaned DEPs ranked by abs(log₂FC). See companion .xlsx for full content.*

##### Table S2. Cleaned DEP set with directional split

Source file: tables/Table_S2_DEPs_cleaned.xlsx (3 sheets: all_DEPs_p_lt_0.05 n = 473; UP_in_LOU n = 308; DOWN_in_LOU n = 165).

**Preview: top 15 DOWN-in-LOU DEPs by |log₂FC|.**

| UNIPROT | GeneSymbol | log2FC | P_value | q_value_BH |
|---|---|---|---|---|
| **O35821** | Mybbp1a | -0.185 | 0.0493 | 0.778 |
| **D4ACN6** | Cert1 | -0.19 | 0.0429 | 0.773 |
| **P10868** | Gamt | -0.193 | 0.0388 | 0.768 |
| **A0A8I5ZKP5** | Ttyh3 | -0.198 | 0.0454 | 0.778 |
| **F7F1X7** | Pmm2 | -0.204 | 0.0308 | 0.751 |
| **Q3KRE0** | Atad3a | -0.214 | 0.032 | 0.751 |
| **Q6AYD9** | Nudt19 | -0.215 | 0.0392 | 0.769 |
| **P12007** | Ivd | -0.22 | 0.0488 | 0.778 |
| **A0A8I6AG33** | Tnpo1 | -0.22 | 0.0427 | 0.773 |
| **P21775** | Acaa1a | -0.22 | 0.0354 | 0.757 |
| **Q5XIG4** | Ociad1 | -0.227 | 0.024 | 0.74 |
| **A0A0G2K3R1** | Map3k4 | -0.228 | 0.0277 | 0.751 |
| **Q6I7R3** | Isoc1 | -0.229 | 0.0477 | 0.778 |
| **Q6P6Q9** | Micu1 | -0.229 | 0.0306 | 0.751 |
| **A0A8I6ALA6** | Frmpd4 | -0.231 | 0.0361 | 0.763 |

**Preview: top 15 UP-in-LOU DEPs by |log₂FC|.**

| UNIPROT | GeneSymbol | log2FC | P_value | q_value_BH |
|---|---|---|---|---|

| | | | | |
|---|---|---|---|---|
| **Q5QE79** | Aox4 | 2.87 | 0.0024 | 0.524 |
| **P62839** | Ube2d2 | 2.82 | 0.0345 | 0.751 |
| **A0A8I6GJU0** | Rnpep | 2.72 | 0.000637 | 0.524 |
| **M0R3X7** | Hkdc1 | 2.56 | 0.0099 | 0.644 |
| **P20759** | nan | 2.01 | 0.0119 | 0.654 |
| **F8WGA3** | Serpinb1b | 1.53 | 0.018 | 0.715 |
| **A0A8I6GMB0** | Pdk1 | 1.39 | 0.0165 | 0.696 |
| **O55207** | Synj2 | 1.38 | 0.0182 | 0.715 |
| **Q5XIF2** | Uros | 1.37 | 0.00636 | 0.607 |
| **D3ZLC4** | nan | 1.27 | 0.00923 | 0.644 |
| **A0A8I6A3C3** | nan | 1.27 | 0.00612 | 0.607 |
| **A0A0G2K3Z7** | Taf15 | 1.25 | 0.0237 | 0.74 |
| **A0A8I6AC14** | nan | 1.25 | 0.00352 | 0.548 |
| **A0A8I6ABN7** | nan | 1.2 | 0.045 | 0.778 |
| **Q5I0K3** | Clybl | 1.2 | 0.0321 | 0.751 |

**Table S3. Over-representation analysis (LOU side)**

Source file: tables/Table_S3_ORA_enrichr.xlsx (10 sheets, one per library × direction: KEGG_2021_Human × {up_in_LOU, down_in_LOU}; same for GO_Biological_Process_2023, Reactome_2022, MSigDB_Hallmark_2020, WikiPathway_2023_Human; total enriched terms across all libraries shown).

**Preview: top 12 KEGG terms enriched on the 165 DOWN-in-LOU DEPs (sorted by P-value).**

| Term | P-value | Adjusted P-value | Genes |
|---|---|---|---|
| **Oxidative phosphorylation** | 0.00755 | 0.315 | NDUFA9;NDUFA13;NDUFB10;NDUFA5;NDUFS1;UQCRC2 |
| **Non-alcoholic fatty liver disease** | 0.01 | 0.315 | NDUFA9;NDUFA13;NDUFA5;NDUFB10;NDUFS1;UQCRC2 |
| **Hypertrophic cardiomyopathy** | 0.013 | 0.315 | ACE;ITGB5;ITGA6;EMD |
| **Diabetic cardiomyopathy** | 0.0168 | 0.315 | NDUFA9;NDUFA13;ACE;NDUFA5;NDUFB10;NDUFS1;UQCRC2 |
| **Fructose and mannose metabolism** | 0.0174 | 0.315 | TKFC;PMM1;PMM2 |
| **Retinol metabolism** | 0.0183 | 0.315 | RETSAT;ALDH1A1 |
| **ECM-receptor interaction** | 0.0193 | 0.315 | ITGB5;COL6A3;ITGA6 |
| **Arginine and proline metabolism** | 0.0213 | 0.315 | GAMT;PRODH;L3HYPDH |
| **Lysine degradation** | 0.0213 | 0.315 | GCDH;TMLHE;PLOD3 |
| **Amyotrophic lateral sclerosis** | 0.0296 | 0.368 | NDUFA9;NUP214;NDUFA13;NDUFA5;NDUFB10;NUP35;NDUFS1;NUP54;UQCRC2;NUP160 |
| **Terpenoid backbone biosynthesis** | 0.0304 | 0.368 | FNTB;HMGCS2 |
| **Thermogenesis** | 0.0368 | 0.391 | NDUFA9;NDUFA13;KDM3B;NDUFA5;NDUFB10;NDUFS1;UQCRC2 |

**Preview: top 12 KEGG terms enriched on the 308 UP-in-LOU DEPs (sorted by P-value).**

| Term | P-value | Adjusted P-value | Genes |
|---|---|---|---|
| **Aminoacyl-tRNA biosynthesis** | 0.00148 | 0.284 | LARS2;MTFMT;TARS3;QRSL1;VARS2;YARS2;CARS2 |
| **Amino sugar and nucleotide sugar metabolism** | 0.00554 | 0.529 | GMPPB;CYB5R4;NAGK;HKDC1;MPI;UAP1 |
| **Collecting duct acid secretion** | 0.0116 | 0.741 | ATP6V1G2;ATP6V1E1;ATP6V1C1 |
| **Glycolysis / Gluconeogenesis** | 0.831 | 0.999 | HKDC1 |
| **TNF signaling pathway** | 0.838 | 0.999 | NFKBIA |
| **RNA degradation** | 0.838 | 0.999 | LSM8 |

| | | | |
|---|---|---|---|
| **Chronic myeloid leukemia** | 0.851 | 0.999 | NFKBIA |
| **Glioma** | 0.851 | 0.999 | PTEN |
| **Osteoclast differentiation** | 0.851 | 0.999 | NFKBIA |
| **Pathogenic Escherichia coli infection** | 0.853 | 0.999 | NFKBIA;MYO5A;MYH10 |
| **Epstein-Barr virus infection** | 0.855 | 0.999 | NFKBIA;APAF1 |
| **Toxoplasmosis** | 0.857 | 0.999 | NFKBIA |

**Table S3b. Over-representation analysis (human AD side)**

Source file: tables/Table_S3b_ORA_human_AD.xlsx (10 sheets, one per library × {B4, B6} × {up, down}; identical settings to Table S3, applied to our published human B4-vs-CT and B6-vs-CT DEG lists from Badina et al., 2025).

**Preview: top 12 KEGG terms enriched on LAD-down DEGs (Braak 6 vs CT).**

| Term | P-value | Adjusted P-value | Genes |
|---|---|---|---|
| **GABAergic synapse** | 5.74e-06 | 0.00137 | NSF;GABRA1;GABARAPL1;SLC12A5;GABRA4;GAD1;GABRA3;CACNA1B;GAD2;GPHN;GABRG2;GNAI1;GLS;GNG3;PRKACB;GABRD |
| **Oxidative phosphorylation** | 9.99e-05 | 0.00676 | ATP6V1G2;ATP5F1B;ATP5PB;NDUFA10;ATP6V1B2;ATP5MC3;ATP6V1H;ATP6V1E1;ATP6V1D;ATP6V1C1 |
| **Synaptic vesicle cycle** | 0.000104 | 0.00676 | NSF;SNAP25;RAB3A;ATP6V1G2;SYT1;STXBP1;CACNA1B;DNM1;DNM3;ATP6V1B2;ATP6V1H;ATP6V1E1;ATP6V1D;ATP6V1C1 |
| **Parkinson disease** | 0.0015 | 0.00676 | ATP5PB;CAMK2A;NDUFA10;ITPR1;ATP5MC3;GNAI1;MAPK10;PSMA5;MAPK8;UCHL1;ATP5F1B;CALM3;VDAC1;CALM1;CALM2;PRKACB;SLC25A4 |
| **Pathways of neurodegeneration** | 0.000154 | 0.00676 | GRIA1;APP;GRIA2;CAMK2A;NDUFA10;CACNA1B;ITPR1;ATP5MC3;GRIN2A;MAPK8;PPP3CB;PPP3R1;UCHL1;ATP5F1B;NEFL;GRIA3;GABARAPL1;NOS2;ATP5PB;SOD1;MAPK10;PSMA5;IL1A;IL1B;CALM3;VDAC1;CALM1;CALM2;SLC25A4 |
| **Nicotine addiction** | 0.0017 | 0.00676 | GRIA1;CHRNB2;GRIA2;GABRA1;GRIN2A;GABRA4;GABRA3;CACNA1B;GABRD;GRIA3;GABRG2 |
| **Retrograde endocannabinoid signaling** | 0.000378 | 0.0123 | GRIA1;GRIA2;GABRA1;GABRA4;NDUFA10;GABRA3;CACNA1B;ITPR1;GABRG2;GNAI1;MAPK10;GNG3;MAPK8;CNR1;PRKACB;GABRD;GRIA3 |
| **Long-term potentiation** | 0.000445 | 0.0123 | GRIA1;GRIA2;PPP3CB;GRIN2A;PPP3R1;CAMK4;CAMK2A;ITPR1;CALM3;CALM1;PRKACB;CALM2 |
| **Calcium signaling pathway** | 0.000467 | 0.0123 | RET;NOS2;PDE1A;CAMK2A;CACNA1B;ITPR1;ATP2B3;ATP2B2;SLC8A1;GRIN2A;PPP3CB;PPP3R1;ADORA2B;CAMK4;PTK2B;CALM3;VDAC1;CALM1;CALM2;PRKACB;SLC25A4 |
| **Circadian entrainment** | 0.000513 | 0.0123 | GRIA1;GRIA2;CAMK2A;ITPR1;GNAI1;GRIN2A;GNG3;NOS1AP;CALM3;CALM1;PRKACB;CALM2;GRIA3 |
| **Dopaminergic synapse** | 0.000861 | 0.0187 | GRIA1;GRIA2;CAMK2A;CACNA1B;ITPR1;GNAI1;PPP2CA;MAPK10;GRIN2A;PPP3CB;GNG3;MAPK8;CALM3;CALM1;PRKACB;CALM2;GRIA3 |
| **Aldosterone synthesis and** | 0.00121 | 0.0239 | CAMK4;CAMK2A;ITPR1;ATP2B3;CALM3;ATP2B2;ATP1A1;ATP1B1;CALM1;PRKACB;CALM2 |

| |
|---|
| **secretion** |

**Table S4. Pre-ranked GSEA outputs**

Source file: tables/Table_S4_GSEA_prerank.xlsx (2 sheets: KEGG_2021_Human n = 306; MSigDB_Hallmark_2020 n = 50). Pre-ranked GSEA on the full hippocampal proteome ranked by $-\log_{10}(p)\cdot \mathrm{sign}(\log_2 FC)$; the mt-Nd2 outlier was excluded from the ranked vector.

**Preview: top 15 KEGG GSEA terms (sorted by FDR q-value).**

| Term | ES | NES | NOM p-val | FDR q-val | Lead_genes |
|---|---|---|---|---|---|
| **Non-alcoholic fatty liver disease** | -0.524 | -2.23 | 0.0005 | 0.0205 | NDUFA13;NDUFA5;NDUFS1;UQCRC2;NDUFA9;NDUFB10;NDUFS4;NDUFS7;NDUFA6;NDUFV1;NDUFA10;UQCR10;NDUFA12;NDUFB3;NDUFS2;NDUFB9;NDUFA2;COX7A2L;NDUFB7;UQCRFS1;NDUFA7;NDUFA4;COX4I1;NDUFC2;COX6B1;UQCRC1;NDUFB8;CYC1;COX7A2;UQCRQ;NDUFV3;NDUFB6;PKLR;UQCRB;PIK3R1;COX5A;NDUFV2 |
| **Oxidative phosphorylation** | -0.497 | -2.08 | 0.0005 | 0.0488 | NDUFA13;NDUFA5;NDUFS1;UQCRC2;NDUFA9;NDUFB10;NDUFS4;NDUFS7;NDUFA6;NDUFV1;NDUFA10;UQCR10;NDUFA12;NDUFB3;NDUFS2;NDUFB9;NDUFA2;COX7A2L;NDUFB7;UQCRFS1;NDUFA7;NDUFA4;COX4I1;NDUFC2;COX6B1;UQCRC1;NDUFB8;CYC1;COX7A2;UQCRQ;NDUFV3;NDUFB6;UQCRB;COX15;COX5A;NDUFV2 |
| **Diabetic cardiomyopathy** | -0.442 | -1.97 | 0.0005 | 0.0128 | NDUFA13;NDUFA5;NDUFS1;UQCRC2;NDUFA9;ACE;NDUFB10;SLC25A5;NDUFS4;NDUFS7;NDUFA6;NDUFV1;NDUFA10;UQCR10;NDUFA12;NDUFB3;NDUFS2;NDUFB9;NDUFA2;COX7A2L;NDUFB7;UQCRFS1;NDUFA7;NDUFA4;MPC2;COX4I1;PLCB3;NDUFC2;COX6B1;UQCRC1;NDUFB8;CYC1;G6PD;COX7A2;UQCRQ;NDUFV3;NDUFB6;UQCRB;SLC25A4;PIK3R1;COX5A;NDUFV2 |
| **Retinol metabolism** | -0.761 | -1.89 | 0.0072 | 0.0244 | RETSAT;ALDH1A1;RDH13;ALDH1A2;AOX1;BCO1 |
| **Aminoacyl-tRNA biosynthesis** | 0.595 | 1.81 | 0.0005 | 0.0365 | YARS2;TARS3;CARS2;LARS2;FARSB;MTFMT;VARS2;CARS1;QRSL1;WARS2;HARS1;FARS2;DARS2;NARS2;AARS1;DARS1;MARS1;FARSA;GATC;MARS2;NARS1;YARS1;LARS1;PARS2;EPRS1;TARS2;SARS1 |
| **ECM-receptor interaction** | -0.549 | -1.83 | 0.0569 | 0.0375 | ITGA6;COL6A3;ITGB5;ITGA3;FN1;TNC;THBS1;ITGAV;DAG1;LAMC1;GP5;ITGB8;AGRN;ITGA7 |
| **mTOR signaling pathway** | 0.538 | 1.83 | 0.0005 | 0.0491 | LAMTOR3;PTEN;ATP6V1E1;MIOS;ATP6V1C1;ATP6V1G2;DEPDC5;ATP6V1D;ATP6V1B1;RPTOR;NPRL3;ATP6V1H;SLC7A5;TSC2;MAPK1;CASTOR2;RRAGB;MLST8;GRB2;WDR59;RRAGA;GSK3B;RPS6KA1;AKT2;RHOA;SLC3A2;PRKAA2;MAPK3;LPIN2;ATP6V1B2;PRKAA1;RAF1;KRAS;PRKCG;PIK3R2;SOS1;PRKCA |
| **Thermog** | -0 | -1 | 0.0 | 0.0 | NDUFA13;NDUFA5;NDUFS1;UQCRC2;NDUFA9;KDM3B;NDUFB10;NDUFS4;NDUFS7;NDUFA6;NDUFV1;NDUFAF6;NDUFA10;UQCR10;NDUFA12;NDUFB3;NDUFS2;NDUFB9;NDUFA2;COX7A2L;NDUFB7;UQCRFS1;ND |

| | | | | | |
|---|---|---|---|---|---|
| **enesis** | .389 | .75 | 005 | 673 | UFA7;NDUFA4;COX4I1;NDUFC2;COX6B1;UQCRC1;NDUFB8;CYC1;COX7A2;UQCRQ;SMARCE1;NDUFV3;NDUFB6;NRAS;RPS6KA3;ACSL3;UQCRB;ACTL6A;COX15;COX5A;NDUFV2;ACSL6;MGLL |
| **Thiamine metabolism** | 0.812 | 1.75 | 0.00151 | 0.0689 | AK5;NTPCR;AK1;ACP1 |
| **Retrograde endocannabinoid signaling** | -0.398 | -1.72 | 0.0005 | 0.0697 | NDUFA13;NDUFA5;NDUFS1;NDUFA9;NDUFB10;NDUFS4;NDUFS7;NDUFA6;NDUFV1;NDUFA10;NDUFA12;NDUFB3;NDUFS2;NDUFB9;NDUFA2;NDUFB7;NDUFA7;NDUFA4;PLCB3;NDUFC2;PTGS2;NDUFB8 |
| **Hematopoietic cell lineage** | -0.594 | -1.73 | 0.00331 | 0.0705 | ITGA6;ITGA3;TFRC;GYPA;MME;CD9;GP5;CD38 |
| **Rheumatoid arthritis** | 0.698 | 1.72 | 0.00289 | 0.0869 | ATP6V1E1;ATP6V1C1;ATP6V1G2;ATP6V1D;ATP6V1B1;ATP6V1H;ATP6AP1;ATP6V1B2;ATP6V0A1;ATP6V0D1;ATP6V0C |
| **Neurotrophin signaling pathway** | 0.495 | 1.66 | 0.000604 | 0.135 | NFKBIA;RAPGEF1;CAMK2B;CAMK2A;MAP3K5;TRAF6;MAPK1;MAPK9;RAC1;GRB2;GSK3B;RPS6KA1;MAGED1;PLCG1;AKT2;RHOA;CAMK2G;ABL1;MAPK7;MAPK3;MATK;CDC42;MAPK12;RAF1;KRAS;CAMK2D;SHC2;NTRK3;KIDINS220;PIK3R2;SOS1;BAD |
| **Synaptic vesicle cycle** | 0.53 | 1.67 | 0.00126 | 0.138 | NSF;ATP6V1E1;ATP6V1C1;ATP6V1G2;DNM1;ATP6V1D;ATP6V1B1;ATP6V1H;NAPA;DNM3;SNAP25;SLC17A6;SLC32A1;ATP6V1B2;ATP6V0A1;SYT1;AP2A2;SLC17A7;ATP6V0D1;CLTC;RIMS1;ATP6V0C;STX1B;VAMP2;SLC6A7;STX1A |
| **Prion disease** | -0.335 | -1.56 | 0.0005 | 0.146 | NDUFA13;NDUFA5;NDUFS1;UQCRC2;NDUFA9;NDUFB10;SLC25A5;NDUFS4;NDUFS7;NDUFA6;NDUFV1;NDUFA10;UQCR10;NDUFA12;NDUFB3;NDUFS2;NDUFB9;NDUFA2;COX7A2L;NDUFB7;UQCRFS1;NDUFA7;NDUFA4;COX4I1;KLC4;MCU;NDUFC2;COX6B1;UQCRC1;NDUFB8;CYC1;C8B;GRIN2A;COX7A2;UQCRQ;NDUFV3;NDUFB6;UQCRB;SLC25A4;LAMC1;PIK3R1;HSPA2;COX5A;NDUFV2 |

**Table S5. Cross-species panel overlay**

Source file: tables/Table_S5_cross_species_overlay.xlsx (4 sheets: per_gene_overlay n = 447; spearman_summary n = 28 curated axes; sign_test_summary n = 28; axis_membership n = 44).

**Spearman correlation summary (LOU $log_2$FC vs human B4/B6 $log_2$FC, per curated axis).**

| contrast | subset | n | spearman_rho | p |
|---|---|---|---|---|
| **B4vsCT** | all_panel_overlap | 37 | 0.201 | 0.233 |
| **B4vsCT** | LOU_sig_only | 2 | | |
| **B4vsCT** | human_sig_only | 37 | 0.201 | 0.233 |
| **B4vsCT** | both_sig | 2 | | |
| **B6vsCT** | all_panel_overlap | 162 | -0.157 | 0.0457 |

| | | | | |
|---|---|---|---|---|
| **B6vsCT** | LOU_sig_only | 17 | -0.211 | 0.416 |
| **B6vsCT** | human_sig_only | 162 | -0.157 | 0.0457 |
| **B6vsCT** | both_sig | 17 | -0.211 | 0.416 |
| **B4vsCT** | axis_Synaptic_inhibitory | 3 | | |
| **B6vsCT** | axis_Synaptic_inhibitory | 7 | -0.143 | 0.76 |
| **B4vsCT** | axis_Synaptic_vesicular | 0 | | |
| **B6vsCT** | axis_Synaptic_vesicular | 4 | | |
| **B4vsCT** | axis_Synaptic_general | 2 | | |
| **B6vsCT** | axis_Synaptic_general | 10 | 0.459 | 0.182 |
| **B4vsCT** | axis_Oligo_myelin | 6 | 0.464 | 0.354 |
| **B6vsCT** | axis_Oligo_myelin | 0 | | |
| **B4vsCT** | axis_OPC | 1 | | |
| **B6vsCT** | axis_OPC | 1 | | |
| **B4vsCT** | axis_Astrocytic | 3 | | |
| **B6vsCT** | axis_Astrocytic | 4 | | |
| **B4vsCT** | axis_Microglia_complement | 2 | | |
| **B6vsCT** | axis_Microglia_complement | 2 | | |
| **B4vsCT** | axis_Mito_OXPHOS | 1 | | |
| **B6vsCT** | axis_Mito_OXPHOS | 1 | | |
| **B4vsCT** | axis_Immune_dont_eat_me | 0 | | |
| **B6vsCT** | axis_Immune_dont_eat_me | 1 | | |
| **B4vsCT** | axis_Stress_proteostasis | 0 | | |
| **B6vsCT** | axis_Stress_proteostasis | 0 | | |

**Binomial sign-test summary against the 0.5 null (per curated axis).**

| **contrast** | **subset** | **n** | **same_sign** | **frac_same** | **p** |
|---|---|---|---|---|---|
| **B4vsCT** | all_panel | 37 | 20 | 0.541 | 0.743 |
| **B4vsCT** | LOU_sig | 2 | | | |
| **B4vsCT** | human_sig | 37 | 20 | 0.541 | 0.743 |
| **B4vsCT** | both_sig | 2 | | | |
| **B4vsCT** | axis_Synaptic_inhibitory | 3 | | | |
| **B4vsCT** | axis_Synaptic_vesicular | 0 | | | |
| **B4vsCT** | axis_Synaptic_general | 2 | | | |
| **B4vsCT** | axis_Oligo_myelin | 6 | 6 | 1 | 0.0312 |
| **B4vsCT** | axis_OPC | 1 | | | |
| **B4vsCT** | axis_Astrocytic | 3 | | | |
| **B4vsCT** | axis_Microglia_complement | 2 | | | |
| **B4vsCT** | axis_Mito_OXPHOS | 1 | | | |
| **B4vsCT** | axis_Immune_dont_eat_me | 0 | | | |
| **B4vsCT** | axis_Stress_proteostasis | 0 | | | |
| **B6vsCT** | all_panel | 162 | 44 | 0.272 | 5.31e-09 |
| **B6vsCT** | LOU_sig | 17 | 4 | 0.235 | 0.049 |
| **B6vsCT** | human_sig | 162 | 44 | 0.272 | 5.31e-09 |
| **B6vsCT** | both_sig | 17 | 4 | 0.235 | 0.049 |
| **B6vsCT** | axis_Synaptic_inhibitory | 7 | 2 | 0.286 | 0.453 |
| **B6vsCT** | axis_Synaptic_vesicular | 4 | 0 | 0 | 0.125 |
| **B6vsCT** | axis_Synaptic_general | 10 | 5 | 0.5 | 1 |
| **B6vsCT** | axis_Oligo_myelin | 0 | | | |
| **B6vsCT** | axis_OPC | 1 | | | |
| **B6vsCT** | axis_Astrocytic | 4 | 2 | 0.5 | 1 |
| **B6vsCT** | axis_Microglia_complement | 2 | | | |
| **B6vsCT** | axis_Mito_OXPHOS | 1 | | | |
| **B6vsCT** | axis_Immune_dont_eat_me | 1 | | | |
| **B6vsCT** | axis_Stress_proteostasis | 0 | | | |
